# Unraveling proteomic chaos by independent component analysis - ClpX proficiency promotes the iron and oxygen limitation responses of *Staphylococcus aureus* and affects the intracellular bacterial behavior

**DOI:** 10.1101/2025.09.04.674172

**Authors:** Larissa M. Busch, Hannes Wolfgramm, Supradipta De, Christian Hentschker, Manuela Gesell Salazar, Meike Kröber, Celina Hopp, Marie-Sofie Illenseher, Alexander Ganske, Stephan Michalik, Alexander Reder, Sven Hammerschmidt, Dorte Frees, Ulf Gerth, Kristin Surmann, Ulrike Mäder, Uwe Völker

## Abstract

In the opportunistic pathogen *Staphylococcus aureus*, protein homeostasis is largely mediated by the Caseinolytic protease (Clp) system. The proteases ClpXP and ClpCP are crucial for general and targeted proteolysis, which rely on the unfoldases ClpX and ClpC interacting with specific targets. However, the global effect on the proteome especially under infection-relevant stresses is not well-understood. To assess the effect of ClpX deficiency during infection-related processes, mass spectrometry-based global proteome profiles of *S. aureus* HG001 wild-type, an isogenic Δ*clpX* mutant, and a *clpX* complemented strain were recorded under control conditions as well as iron and oxygen limitation. The proteomic profiles revealed specific ClpX- and stress-dependent changes. A set of 24 robust stress-independent ClpX modulated proteins was identified and the stress-dependent influences were unraveled by independent component analysis (using the iModulon approach). These analyses revealed a role of ClpX in e.g., cell division, cell envelope homeostasis, the quinone stress response and prophage activation. Moreover, ClpX-dependent stress-specific effects were observed in the Δ*clpX* mutant, e.g. reduced induction of the heme uptake system under iron limitation and a dampened Rex-controlled oxygen limitation response. This revealed in particular that ClpX is central for heme homeostasis in *S. aureus*. Furthermore, in a *Galleria* infection model, the *S. aureus* Δ*clpX* mutant was attenuated compared to the wild-type HG001. This is consistent with a drastically reduced intracellular replication of the Δ*clpX*-mutant in cell culture-based infection experiments, however, high intracellular persistence of the Δ*clpX* mutant was also observed. This highlights the relevance of ClpX for bacterial fitness and virulence.

**Importance:** During infection processes, pathogens cope with host-mediated stressors. In response to those stressors, bacteria adapt their gene expression as well as their proteome profile. In the pathogen *Staphylococcus aureus*, protein homeostasis is mainly controlled by the Clp system. In particular, ClpX is the most conserved Clp unfoldase and is involved in overall regulation of virulence and bacterial fitness. However, the majority of ClpX targets remains elusive in *S. aureus*. With our proteomics approach and in depth data analysis, we provide a resource for global insight into ClpX-dependent adaptation of *S. aureus* physiology under infection-relevant conditions. Based on this, we uncover ClpX’s role as a central player in the iron and oxygen limitation response. In addition, we demonstrate the importance of ClpX in *S. aureus* bacterial fitness in infection processes. However, reduced levels of ClpX lead to high intracellular persistence, which questions ClpX’s suitability as a therapeutical target.

## Background

Better understanding of the infection processes requires comprehensive knowledge of the interactions between host and pathogen. Consequently, when analyzing bacterial pathophysiology, it is important to consider factors beyond a list of theoretically expressible virulence genes and to analyze how bacteria adapt their proteome to cope the stresses they encounter during infections. The Gram-positive bacterium *Staphylococcus aureus*, an opportunistic pathogen, is associated with more than one million deaths per year (1) and causes severe diseases such as endocarditis, osteomyelitis and sepsis (2). However, it is so called Janus-faced (3), since it also colonizes the skin and anterior nares of healthy human individuals.

The highly conserved caseinolytic protease (Clp) system is crucial for the maintenance of cellular protein homeostasis (4–6). The Clp system in *S. aureus*, consists of the peptidase subunit ClpP and the four AAA+ ATPase chaperones ClpX, ClpC, ClpB and ClpL. Of these, ClpX and ClpC can serve as unfoldase subunits for the Clp protease complex (4, 7). ClpP protein subunits form a barrel-shaped peptidase consisting of two heptameric rings (8). The heptameric ClpP rings form seven hydrophobic pockets, which act as interaction points with the Clp-ATPases, ClpX and ClpC that form hexameric rings (8). The Clp-ATPases interact with ClpP *via* a conserved *h*G[F/L] motif (9). Interaction with adaptor proteins such as the ClpC-specific TrfA (10) or the ClpX-specific YjbH (11) defines a spectrum of specific protease targets. Consequently, the Clp protease system shapes the protein composition of *S. aureus* cells through both specific and general proteolysis.

Despite the description of exemplary target proteins of the Clp system, the exact spectrum of Clp protease and Clp chaperone targets, as well as the physiological consequences of Clp disruption, are still not fully understood. The best-studied staphylococcal Clp protease target are the ClpXP-degraded oxidative and disulfide stress regulator protein Spx and the two cell division hydrolase Sle1 and CxcA (12–14), while chaperone function of ClpX independent of ClpP promotes cell division (15). By influencing protein levels of such transcriptional regulators, the Clp system also induces a pronounced additional and indirect modulation of the cell’s proteomic profile.

During the course of an infection, *S. aureus* encounters typical stresses including limitation of iron or oxygen (16–18). In order to overcome these infection-associated stresses, *S. aureus* mounts both general and specific stress responses. The Clp system and, in particular, the subunit ClpX are implicated in infection-related aspects of *S. aureus* physiology and thus the Clp components are involved in infection-relevant stress responses as those mounted upon oxidative stress (19–21), oxygen limitation (20, 22, 23), heat or cold shock (19, 24), and iron limitation (25, 26). Furthermore, a general role in the modulation of virulence factor levels has especially been demonstrated for ClpP and ClpX (19, 27, 28). Hence, *S. aureus* lacking the ClpXP protease is attenuated in pneumonia and skin infection models and provokes only a reduced host immune response (29, 30).

Given the current lack of understanding of ClpX-driven molecular mechanisms in infection processes and the physiological consequences of ClpX deficiency under infection-relevant stresses, we analyzed the role of ClpX deficiency under infection-relevant conditions by generating global label-free proteome profiles combined with in-depth data analysis. To the best of our knowledge, no state-of-the-art proteome profiles of ClpX-deficient *S. aureus* strains under such stress conditions are available to the community. Accordingly, this analysis provides new insights into ClpX’s role in physiological adaptation to changes related to the niches of *S. aureus* during infection processes and will serve as a valuable resource for further research.

## Material & Methods

### Bacterial strains

Bacterial strains and plasmids used are listed in Table 1, primers used in this study are listed in Supplemental Data Table A.

**Table 1.** Bacterial strains and plasmids.

| *S. aureus* strains | Relevant genotype/ characteristics | Reference |
| --- | --- | --- |
| HG001 | *rsbU*<sup>+</sup>-repaired derivate of NCTC8325 (31) | (32) |
| SLB001 | HG001 $\Delta clpX/clpX::kanR$ | This study |
| SLB002 | HG001 $\Delta clpX$ pTripleTREP_ *clpX* | This study |
| SLB003 | HG001 pJL-sar-GFP<sub>redopt</sub> | This study |
| SLB004 | HG001 $\Delta clpX$ pJL-sar-GFP<sub>redopt</sub> | This study |
| Plasmids |  |  |
| pSauSE | allele exchange plasmid derived from pBASE6 (33) | This study |
| pSauSE_ $\Delta clpX::kanR$ | allele exchange plasmid for replacement of the *clpX* gene against a kanamycin resistance cassette | This study |
| pTripleTREP_ *clpX* | expression plasmid pTripleTREP (34) for controllable *clpX* expression | This study, (34) |
| pJL-sar-GFP<sub>redopt</sub> | pJL74 (35) derivate expressing staphylococcal codon-optimized *gfp* | This study |

The assembly of the newly constructed plasmids was performed using *E. coli* Stellar^TM^ cells (TaKaRa Bio, Japan) *via* Sequence and Ligase Independent Cloning (SLIC) (36). Plasmids isolated from *E. coli* were passed through *S. aureus* RN4220 (37) before transformation of the respective target strain *via* electroporation (38). Amplifications of fragments for cloning were generated by polymerase chain reaction (PCR) using the High-Fidelity Phusion polymerase (New England Biolabs, Germany).

The allele exchange plasmid pSauSE was derived from pBASE6 (33), the construction of the empty pSauSE plasmid is described in detail in Supplemental Data Method A. Based on the empty pSauSE plasmid, pSauSE_Δ*clpX*::*kanR* was constructed. Allele exchange was performed with minor adaptations according to the procedure previously described (39). Briefly, for plasmid integration at the *clpX* locus the strain was cultivated in two passages at 43 °C overnight in tryptic soy broth (TSB; BD, USA) with 10 µg/ml chloramphenicol (Sigma-Aldrich, USA) and one passage for colony separation on TSB agar with 10 µg/ml chloramphenicol. SaeRS functionality after heat incubation was verified by hemolysis activity at 37 °C on Colombia agar plates containing 5% sheep blood as mutation in the *sae*-locus is a common site effect of temperature-dependent mutagenesis in *S. aureus* (40). Then, plasmid excision was induced by cultivation at 35 °C. A relatively high non-permissive temperature of 35 °C for plasmid replication was chosen to avoid selection of low temperature-induced Δ*clpX* suppressor mutants (41). Counter-selection of plasmid-carrying cells was performed by 0.5 µg/ml anhydrotetracycline (AHT; Sigma-Aldrich, USA). Subsequently, mutant candidates were selected based on the antibiotic resistance profile against 10 µg/ml chloramphenicol (negative), 0.5 µg/ml AHT (positive), and 7 µg/ml kanamycin (positive) on TSB agar replica plates.

The complete *clpX* coding sequence was replaced with the kanamycin resistance cassette gene (Figure S1) coding for the aminoglycoside O-phosphotransferase APH(3’)-IIIa (GeneBank: EHC3224956.1) and the resulting mutant strain HG001 Δ*clpX* (SLB001) was verified by PCR, DNA sequencing and Northern Blot analysis.

The *clpX* expression plasmid pTripleTREP_*clpX* was constructed based on the controllable expression plasmid pTripleTREP (34). The *clpX* gene including its transcription start site and the native *clpX* terminator were inserted into the pTripleTREP vector using the primers indicated in Supplemental Data Table A. To construct the complemented strain HG001 *clpX*compl (SLB002), the deletion strain HG001 Δ*clpX* was transformed using the expression plasmid pTripleTREP_*clpX* (Figure S1). The strain was verified *via* PCR, DNA sequencing and Northern Blot analysis. The GFP-labelled strains (SLB003 and SLB004) were generated by transformation of HG001 and the Δ*clpX* mutant using the pJL-sar-GFP_redopt_ plasmid. Construction of pJL-sar-GFP_redopt_ is described in Supplemental Data Method B.

### Cultivation of bacteria

Bacteria were cultivated in TSB supplemented with 20 ng/ml AHT to induce *clpX* expression in the complemented strain, as previously optimized to obtain approximately the same ClpX protein levels as in the HG001 wild-type (34). The *S. aureus* main TSB cultures were inoculated with an exponentially growing TSB-overnight culture to an optical density at 540 nm (OD_540_) of 0.05. They were grown aerobically at 37 °C and orbital shaking at 220 rpm in an air Incubator Shaker Innova 44 (New Brunswick Scientific, USA). The complemented strain HG001 *clpX*cmpl was precultured overnight in TSB supplemented with 10 µg/ml chloramphenicol. In general, overnight precultures did not contain AHT. Besides growth under control conditions, bacteria were also subjected to iron limitation or oxygen limitation. Iron depletion was achieved by addition of the divalent metal ion chelator 2,2’-dipyridyl (DP; Sigma-Aldrich, USA) at a concentration of 600 µM, followed by incubation of the medium at 37 °C for at least 1 h. This method is widely used to introduce iron limitation and iron is shown to be the main growth prohibiting factor in DP treated TSB (16, 42). Oxygen limitation was achieved by using the maximal volume of the cultivation flask (43). Bacteria were assessed in exponential (2.5h after inoculation of the main culture – ranging from OD_540nm_ ∼0.5 to ∼2.0 depending on the stress condition and the cultivated strain) and stationary growth phase (8h after inoculation of the main culture – ranging from OD_540nm_ ∼5 to ∼15) to gain insight into bacterial cells in different growth states reflecting also different states of infection processes (44, 45).

For infection experiments, bacterial strains were cultured aerobically at 37 °C in prokaryotic minimal essential medium (pMEM: 1x MEM (Biochrom AG, Germany), 1x non-essential amino acids (PAN-Biotech GmbH, Germany), 4 mM L-glutamine (PAN-Biotech GmbH, Germany), 10 mM 2-[4-(2-hydroxyethyl)piperazin-1-yl]ethanesulfonic acid (HEPES; PAN-Biotech GmbH, Germany), each 2 mM alanine, valine, leucine, isoleucine, aspartate, glutamate, serine, threonine, cysteine, proline, phenylalanine, histidine and tryptophan (Sigma-Aldrich, USA); sterile filtrated; pH 7.4) as described (46). Precultures contained 10 µg/ml erythromycin (Sigma-Aldrich, USA) and 0.01% (w/v) yeast extract (VWR, Germany). Exponentially growing samples cultivated in pMEM were used for inoculation of the main culture to an optical density at 600 nm (OD_600_) of 0.05.

### Measurements of pH during growth of bacterial cells

During cultivation of the bacterial strains in TSB with the applied infection-relevant stresses as described above, pH of the culture was measured with pH indicator stripes (pH 5.0-10.0 MQuant Merck, Germany; pH 4.0-7.0 MQuant, Merck, Germany; pH 6.0-7.7 pH-Fix, Macherey-Nagel, USA; pH 7.5-14.0 Alkalit, Merck, Germany). At least two different types of stripes were used per measurement.

### Cell culture infection experiments with 16HBE14o-cells

For infection experiments, the human epithelial cell line 16HBE14o-, a transformed bronchial epithelial cell line (47), was employed as described before (48) and summarized in Supplemental Methods C. Four days prior to infection, cells were seeded into 12-well plates without (cell counting) or with (fluorescence microscopy) sterile high-precision cover slips (18 mm diameter; Roth, Switzerland) or into a 24-well glass bottom plate (live cell imaging; Ibidi, Germany) at a concentration of 1×10^5^ cells/cm^2^.

Internalization experiments were performed as described previously (48) with slight modifications. In brief, 16HBE14o-cells were counted using the automated cell counter Countess (Invitrogen, Germany) according to manufacturer’s instructions and infected at a multiplicity of infection (MOI) of 50 bacteria per host cell (cell counting and fluorescence microscopy) and of 40 (live cell imaging). To prepare the master mix for infection, a mid-exponential (OD_600_ of 0.4) culture of HG001 pJL-sar-GFP_redopt_ or Δ*clpX* pJL-sar-GFP_redopt_ in pMEM was diluted in eMEM and buffered with 2.9 µl sodium hydrogen carbonate (7.5%; PAN-Biotech GmbH, Germany) per ml bacterial culture. The eMEM of the confluent 16HBE14o-cells was aspirated and replaced with the master mix. Bacterial culture as well as the remaining master mix was collected on ice for bacterial counting as described below. The eukaryotic-bacterial co-culture was incubated for 1 h at 37 °C in 5% CO_2_ humidified atmosphere. Subsequently, an aliquot of the supernatant was collected to count the number of non-adherent bacteria. The remaining medium was removed and replaced with eMEM containing 10 µg/ml lysostaphin (AMBI Products LLC, USA). Afterwards, cells and bacteria were co-cultured for one week without further medium exchange. To monitor the course of infection, human host cells and internalized bacterial cells were counted at selected time points post infection. Thus, for each strain and time point, the medium of one well was removed, infected host cells were washed with PBS and detached from the plate with trypsin-EDTA (PAN Biotech, Germany). Epithelial cells were counted from this suspension using a Neubauer counting chamber. The remaining suspension was treated with 1% (w/v) sodium dodecyl sulfate (SDS) to a final concentration of 0.05% (w/v) SDS. Numbers of intracellular bacteria were determined with a Guava® easyCyte flow cytometer (Merck Millipore, Germany) by excitation of the GFP with a 488 nm laser and detection at 525/15 nm.

### Imaging of infection processes

For detailed fluorescence microscopy of selected time points post infection with HG001 pJL-sar-GFP_redopt_ or Δ*clpX* pJL-sar-GFP_redopt_, infected 16HBE14o-cells were treated as described before (49), but without additional staining of the bacteria. In brief, the infected epithelial cells on coverslips were washed with DPBS (PAN Biotech, Germany) and fixated with 2% formaldehyde for a minimum of 2 h at 4 °C. Afterwards, cells were washed twice with PBS and stored in PBS at 4°C until staining. The staining procedure was initiated by permeabilizing cell membranes with 0.1% Triton X-100 in deionized water for 5 min at room temperature. DNA was stained with Hoechst 33258 (1 µg/ml; Thermo Fisher Scientific, USA) and F-actin filaments with Phalloidin conjugated to Alexafluor 568 (1 U/ml; Thermo Fisher Scientific, USA) in one single solution for 10 min at room temperature. The cells on the coverslips were washed twice with PBS and once thoroughly with deionized water to remove salt residues. The coverslips were mounted with 20 µl DAKO fluorescence mounting medium (DAKO, Germany) upside down to an object slide and were allowed to dry for at least 1 h prior to microscopy. Fluorescence microscopy was performed with a Leica DM2500 LED (Leica Microsystems, Germany) microscope, equipped with the objectives HC PL FLUOTAR 10 x and HC PL FLUOTAR L 40 x, a CoolLED pE-300white (SB) light source (CoolLED, UK) and a Leica DFC3000 G camera. Fluorescence signals of GFP of the bacteria (excitation BP 470/40, emission BP 525/50; exposure time 35 ms), Phalloidin conjugated Alexafluor 568 (excitation BP 560/40, emission BP 630/75; exposure time 400 ms) and DNA-labeling Hoechst 33258 (excitation: BP 350/50, emission: BP 460/50; exposure time 150 ms) were detected. LAS X software (v 3.3.3.16958; Leica Microsystems, Germany) was used for primary analysis and visualization. Quantification and image preparation was performed using FIJI (v153q; (50)) with the Bio-Formats plugin (51). Subsequent normalization of the mean GFP intensity to the mean Hoechst 33258 intensity and visualization was performed using R (v4.1.2) with tidyverse (v2.0.0; (52)).

For live cell imaging, infection experiments were performed in 24-well glass bottom plates as described above at a MOI of ∼40. After addition of lysostaphin, plates were subjected to a Leica DMi8 Stellaris 8 microscope (Leica Microsystems, Germany) to a chamber maintaining 37 °C, 5% CO_2_, and a humidified atmosphere. Under these conditions, pictures were acquired every 30 min starting 2 h post infection for one day. Thereby, GFP of the plasmid carrying *S. aureus* cells was detected using the FITC filter cube (Excitation: BP 480/40, Emission: BP 527/30) with an exposure time of 0.01 s. The host cells were recorded in bright field mode. For recording, a HC PL APO 40x/1.10 W CORR CS2 objective and a Hamamatsu Flash 4.0 V3 camera (Hamamatsu Photonics, Japan) were used. LAS X software (v4.6.1.27508) was used for data acquisition. Live cell microscopy data were subsequently analyzed and quantified using FIJI (v153q) with the Bio-Formats plugin. Mean GFP intensity was quantified script-based for each single picture of the whole time series. Subsequently, normalization to the MOI of 40, loess regression-based background-correction (using the time points 2 h, 12.5 h and 25.5 h as anchor points) and visualization were performed in R (v4.1.2) with tidyverse (v2.0.0) using the modelr (v0.1.11) package.

### Galleria mellonella infection model

For the *Galleria mellonella* infection model, the *S. aureus* strains HG001 pJL-sar-GFP_redopt_ and Δ*clpX* pJL-sar-GFP_redopt_ were cultivated in pMEM as described above in three independent experiments. Exponentially growing bacterial cells (10 ml) were harvested at OD_600_ 0.4 by centrifugation (8,000 ×g, 10 min, room temperature (RT)). Bacteria were washed in sterile 0.9% (w/v) NaCl solution and again pelleted by centrifugation (8,000 ×g, 10 min, RT). Bacteria were then resuspended in 2 ml 0.9% (w/v) NaCl solution and the bacterial concentration was determined based on the GFP reporter in the Guava® easyCyte flow cytometer. Subsequently, the bacterial concentration was adjusted to 1×10^5^ bacteria per 10 µl injection volume (1×10^7^ cells/ml). Ten *Galleria mellonella* larvae (proinsects Gmbh, Germany) per biological replicate and strain were intrahemocelically injected at the last-but-one proleg. Per replicate, one mock infection group (0.9% (w/v) NaCl) was included, resulting in a total of 40 larvae per strain or mock infection for each of the three biological experiments. A gastight microliter syringe (Hamilton, USA) coupled with a repeating dispenser (Hamilton, USA) was used to ensure equal infection doses as previously described in (53).

The mock- or *S. aureus*-infected larvae were kept at 37 °C in the dark for ten days. The survival number of the larvae was tracked every 24 h. Survival analysis and visualization were performed in R (v4.1.2) with tidyverse (v2.0.0) using the survival (v3.6.4) and ggsurvfit (v1.1.0) packages. Growth conditions for the *Galleria mellonella* larvae are described in Supplemental Data Method D.

### Sample harvest and preparation for mass spectrometric analysis

*S. aureus* strains HG001, Δ*clpX* and Δ*clpX* pTripleTREP_*clpX* were cultivated in TSB and under infection-relevant stress conditions as described above. Bacterial cells were harvested in exponential and stationary growth phase by centrifugation (4 °C, 10,000 xg, 3 min). Cell pellets were washed in cold 20 mM HEPES (Sigma-Aldrich, USA) and pellets were immediately frozen in liquid nitrogen. For mechanical cell disruption, frozen pellets were suspended in 100 µl 20 mM HEPES + 1% (w/v) SDS per ∼15 OD unites of harvested culture. Then, suspended samples were disrupted in a bead mill (Retsch GmbH, Germany; 3 min, 30 Hz) and the bacterial cell powder was resuspended in 20 mM HEPES + 1% SDS using four-times the volume used in the bead mill. The lysates were treated with Pierce™ Universal Nuclease (Pierce, Thermo Fisher Scientific, USA; 2.5 U, 4 mM MgCl_2_) for 20 min at 37 °C followed by ultra-sonication for 5 min in an ultrasonic bath (Sonorex, Germany). To remove cell debris, samples were centrifuged (30 min, 17,000 xg, RT). Protein concentrations were determined using a Micro BCA^TM^ Protein Assay Kit (Pierce, Thermo Fisher Scientific, USA) and analyzed as described (54). The protein mixtures per sample were spiked with 10% (w/w) of a heavy labelled ^15^N *Bacillus subtilis* standard protein mixture similar to (55) and as described in Supplemental Data Method E.

Tryptic digestion of proteins and peptide purification were performed as recently described (54) with minor adjustments: 100 µg of a 1:1 mixture of hydrophilic (GE Healthcare, UK) and hydrophobic (Thermo Fisher Scientific, USA) carboxylate-modified magnetic SeraMag Speed Beads were added to 5 µg protein mixture per sample (4.5 µg sample proteins + 0.5 µg ^15^N external standard proteins). Samples were incubated for 10 min at RT in 80% (v/v) acetonitrile (ACN). The beads with associated proteins were then washed twice with 80% (v/v) ethanol and once with 100% ACN at a magnetic rack. For protein digestion, the beads were re-buffered into 50mM Tris-HCl 1mM CaCl_2_ (pH 8.0) and incubated with 200 ng Trypsin/LysC Mix (Promega, USA) for 16 h at 37 °C. The digestion was stopped and peptides were eluted by addition of 0.5% (v/v) trifluoracetic acid. After two centrifugation steps (1 min, 17,000 xg, RT) and incubation on the magnetic rack, peptide samples were fully separated from the beads and transferred to an MS vial.

### Mass spectrometric data acquisition

Tryptic peptide solutions were separated for nanoLC-MS/MS analysis on an Ultimate 3000 nano-LC system (Thermo Fisher Scientific, USA) and analyzed in data-independent acquisition (DIA) mode on an Orbitrap Exploris^TM^ 480 mass spectrometer (Thermo Fisher Scientific, USA). For further details, see Supplemental Data Table B and C.

DIA-MS data was analyzed using a spectral library-based approach in the Spectronaut software (version 18.6.231227.55695; Biognosys AG, Switzerland). A dedicated spectral library was built for this project, comprising DIA- and DDA-MS measurements of NCTC 8325 lineage strains under several stress and infection conditions measured in 320 runs on the Orbitrap Exploris^TM^ 480 and the Q Exactive^TM^ HF mass spectrometers (77,137 precursors; 2,427 protein groups).

Database searches were performed against the *Aureo*Wiki *S. aureu*s NCTC 8325 protein database (56) where the RsbU sequence was replaced by the *S. aureus* Newman RsbU sequence to represent the *S. aureus* HG001 protein sequence (2853 staphylococcal protein sequences; three marker protein sequences for TetR, Cat and KanR; four contaminant protein sequences). For normalization, a second spectral library, integrating measurements of the complex heavy labelled *B. subtilis* standard (14,768 precursors), was created by searching against a *B. subtilis* 168 protein data base (4,201 protein sequences).

Detailed parameters for the search and library construction are summarized in Supplemental Data Table D. Ion intensities were global median normalized based on only the heavy labelled ions identified in all samples to allow robust and reliable relative quantification despite large strain- and cultivation condition-related differences in the cellular proteome.

The mass spectrometric proteomics data, corresponding protein data bases and spectral libraries have been deposited to MassIVE (https://massive.ucsd.edu) repository with the dataset identifier MSV000095845 (ProteomeXchange dataset PXD055808).

Global spike-in-normalized Spectronaut-processed data was further analyzed in R (v4.4.1). R packages used for proteome data analysis and visualization are listed in Supplemental Data Table E. Heavy-labelled *B. subtilis* standard proteins and methionine-oxidized peptides were removed from the data set prior to quantitative analyses. Spectronaut-processed data was quality validated and prepared for downstream analysis using an in-house version of the SpectroPipeR pipeline (55, 57).

For global proteome analysis, peptide ions were filtered for ions identified (q-value ≤ 0.001) in at least 50% of one sample condition. Peptide intensities were calculated as sum of corresponding ion intensities. Subsequently, peptide intensities were condition-wise median-median normalized to reduce noise introduced by manual addition of the heavy-labelled standard. Peptide-based ROPECA statistics (58) were calculated protein-wise for single condition comparisons with subsequent p-value adjustment using the Benjamini-Hochberg method (59). Only protein groups identified with at least two peptides were considered for ROPECA statistics and further protein-based analyses. For pairwise comparisons, proteins were only considered if reliably identified in at least one of the compared conditions (at least two peptides with q-value ≤ 0.001 in at least 50% of the condition). Non-unique protein groups were removed.

Based on the normalized peptide intensities, protein levels were estimated using the MaxLFQ algorithm (60). MaxLFQ values were further median normalized and adjusted to the mean relative Spectronaut-based iBAQ (61) values to allow precise estimation of differences between the conditions *via* the MaxLFQ approach and estimation of the general abundance of the particular protein using the iBAQ approach. Non-unique protein groups were removed. Gene set enrichment analyses (GSEA; (62)) were generally performed using the fgsea R-package (v1.20.0; (63)).

### Decomposition analyses

For analysis of protein profiles, adjusted maxLFQ protein levels were used. Principal component analysis was performed using R (v4.1.2) with tidyverse (v2.0.0) and in particular the R package FactoMineR (64) on log_2_-transformed (and half-minimal zero imputed) adjusted maxLFQ protein levels to ensure Gaussian distribution. Further, for each component the separation of proteome profiles according to known condition parameters was tested using the Kruskal-Wallis test. Proteins spanning the components were tested for enrichment of known regulons according to RegPrecise (65)-based *Aureo*Wiki regulons ((56); downloaded February 2025) using the fgsea package (v1.20.0).

For the iModulon-inspired independent component analysis (ICA), the same log_2_-transformed adjusted maxLFQ protein levels were exploited. More detailed information about the methodical background is provided in Supplemental Data Method F. Data was centered to the mean protein value of HG001 wild-type under TSB control condition in the exponential growth phase for each protein (Figure S8). Centered data was visualized using the R-package ComplexHeatmap (v2.15.4; (66)). ICA was performed using the FastICA (67) implementation in the python (v3.7) packages scikit-learn (v1.0.2). To determine the optimal independent components, the OptICA method ((68); --iter 250 --tolerance 1e-7 --min-dim 2 --step-size 5) was exploited in the implementation provided by iModulonMiner (69). Detected independent components, so called i-modulons, were associated with condition parameters using the Kruskal-Wallis test based on the i-modulon activity. Known regulons and sRNA targetomes according to *Aureo*Wiki (56) and (55) as well as a robust ROPECA-based ClpX modulon, as defined in this study, were associated *via* GSEA on the protein loadings per i-modulon.

To define sets of i-modulon members, we exploited the criterion of non-gaussian distribution of protein loadings for each i-modulon, similar to (70). For each i-modulon, the absolute most loaded proteins were excluded one by one and the remaining protein weights were tested for non-gaussian distribution using the D’Agostino omnibus test (71) in the implementation of the fBasics R-package (v4022.94). We used a p-values threshold (Bonferroni-corrected for the number of i-modulons) of 0.001 or, if not reached in the first half of the sorted protein weights, the maximum of the adjusted p-values to consider the excluded proteins as significant i-modulon members.

### Western Blot for validation of ClpX protein levels

Cellular protein extracts prepared as described above for the proteome analysis were used for validation of ClpX protein levels *via* Western Blot analysis as previously described (72). For each sample, 5 µg of protein mixture was analyzed. Proteins were detected using the LI-COR system (LI-COR Biosciences, USA). For total protein detection, the LI-COR Revert Total Protein Stain protocol was used according to the manufacturer’s instructions and for ClpX detection, the primary polyclonal α-ClpX*_B. subtilis_* antiserum (1:5,000, (73), in *Intercept*^®^ T20 AB Diluent, LI-COR Biosciences) and the secondary IRDye 800CW goat-anti-rabbit IgG antibody (1:10,000 in *Intercept*^®^ T20 AB Diluent containing 0.01% (v/v) SDS) were used.

### Northern Blot analysis of IsrR sRNA levels

The *S. aureus* strains HG001 and Δ*clpX* were cultivated under control and iron limitation conditions as described above. In exponential and stationary growth phase, 15 OD_540_ units of bacterial cells were collected, rapidly cooled down in liquid nitrogen and centrifuged (4 °C, 10,000 xg, 3 min). Bacterial pellets were frozen in liquid nitrogen and stored at -70 °C. Subsequent mechanical cell disruption and RNA preparation were carried out as described previously (74). For each sample, 4 µg of total RNA was used for Northern Blot analysis of IsrR levels. Northern Blotting using the IsrR probe was performed as recently described (42).

## Results

### ClpX deficiency and infection-relevant stress conditions affect growth of *S. aureus* HG001

To investigate the effect of ClpX deficiency on the *S. aureus* proteome profile under infection-relevant stresses, we generated a Δ*clpX* mutant (Figure 1A & S1) in the *S. aureus* laboratory strain HG001 (32). For complementation, the controllable expression plasmid pTripleTREP was used to obtain physiological protein levels of ClpX (Figure 1B & S2; (34)).

**Figure 1:**
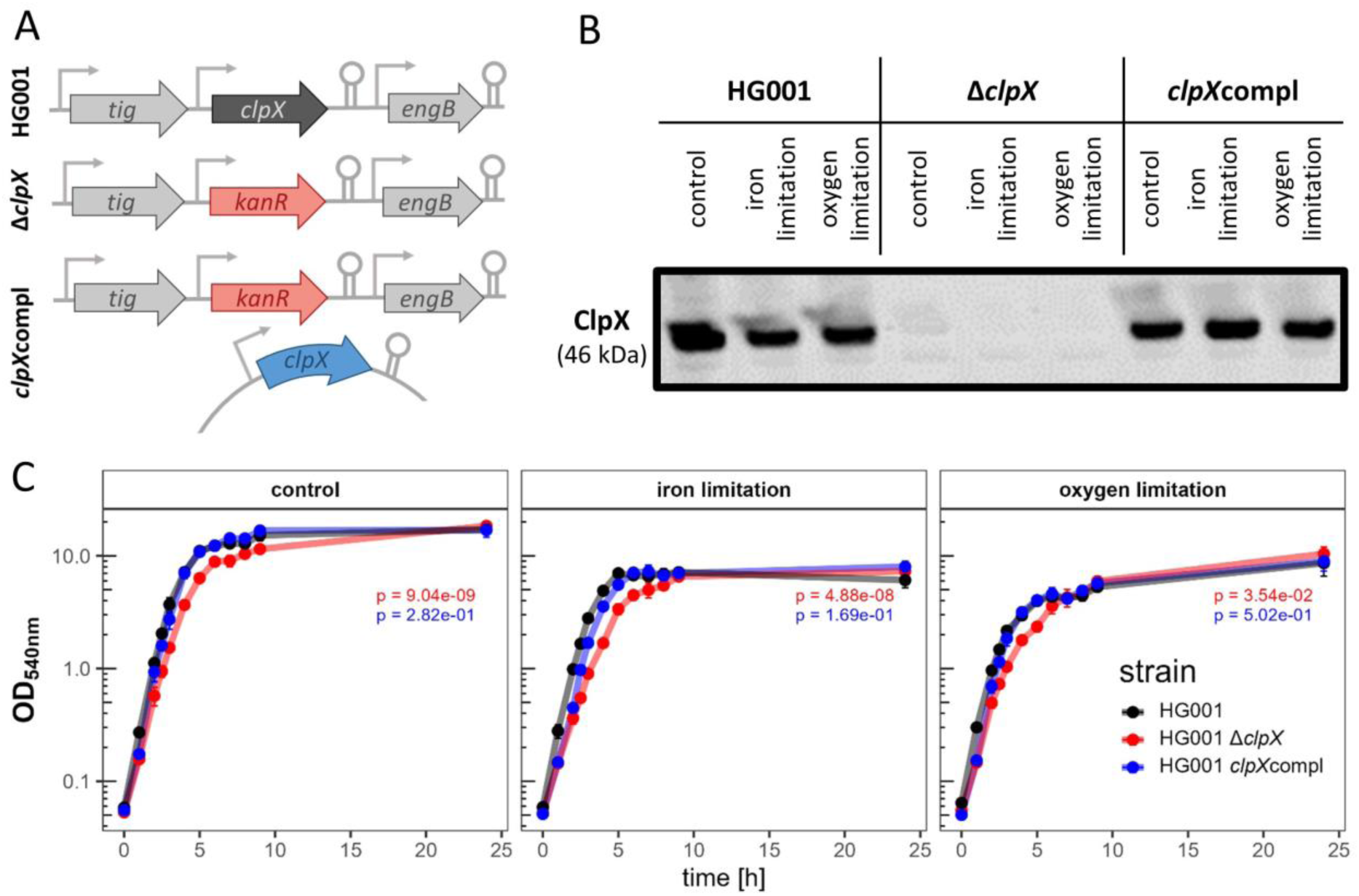
Overview of HG001, Δc*lpX* and *clpX*compl and cultivation of the strains under infection-relevant conditions. (A) Representation of the strains used in this study to investigate the role of ClpX deficiency on the physiology of *S. aureus* under infection-relevant conditions. (B) Western Blot analysis of ClpX for the three strains under infection-relevant conditions. Samples were harvested in exponential growth phase (2.5h). 5 µg of total protein were used. (C) Growth curves. The strains HG001, Δc*lpX* and *clpX*compl were cultivated under control condition (TSB), iron limitation (TSB + 600 µM DP) and oxygen limitation (TSB 100% flask volume). All media contained 20 ng/ml AHT. Four independent biological replicates were cultivated. Standard deviation is indicated as error bars. Differences in general growth was tested by time point-paired t-test between the HG001 wild-type and Δc*lpX* (p-value depicted in red) and between the HG001 wild-type and *clpX*compl (p-value depicted in blue) per cultivation condition.

During infection, *S. aureus* encounters limitations in iron and oxygen (75). To study the effect of these stresses in combination with ClpX deficiency, the HG001 wild-type, the Δ*clpX* mutant and a *clpX* complemented strain were cultivated under iron-and oxygen-limited conditions, respectively (Figure 1C). As expected, iron and oxygen limitation caused earlier entry into the stationary and reduced growth yield of all strains (42, 76). For the HG001 wild-type, the doubling time under control conditions during the exponential growth phase (up to 4 h of cultivation) was increased from 34.6 ± 0.5 min to 37.6 ± 0.9 min under iron limitation and to 43.4 ± 0.3 min under oxygen limitation. Likewise, the final OD_540_ at 24h of cultivation was also reduced (Figure 1C). With regard to ClpX, the growth rate of the Δ*clpX* mutants was reduced under all conditions compared to the ClpX-proficient strains (doubling time for control: 39.3 ± 0.8 min, iron limitation: 47.9 ± 0.4 min; oxygen limitation: 47.8 ± 1.1 min). In agreement with this observation, ClpX was demonstrated to be involved in controlling of the cell wall metabolism and cell division in *S. aureus* (13–15, 41, 77, 78).

### Proteome profiles reveal an interaction between infection-relevant stress conditions and ClpX deficiency

Although a few studies on Clpx-dependent changes of the *S. aureus* proteome have been carried out (e.g. 26, 79), there is a lack of state-of-the-art proteomic profiles of a ClpX-deficient *S. aureus* strain under infection-relevant conditions. To provide such a resource, the cellular proteome profiles of bacterial cells grown under control conditions, iron limitation and oxygen limitation were recorded during exponential and stationary growth and systematically analyzed (Figure S4).

A total of 2,110 protein groups were identified, of which 1,926 staphylococcal proteins and the three marker proteins TetR, Cat and KanR were identified with at least two peptides in single-protein groups. In this manner, 67.51% (1,926/2,853) of the annotated protein encoding genes and 84.09% (1,929/2,294) of the protein groups with at least two peptides in the here provided spectral library were quantified in the presented data set.

A principal component analysis (PCA; Figure 2) was employed to investigate proteome profiles on a global scale. The PCA was calculated using the iBAQ-rescaled maxLFQ protein level estimations (Table S1; Data S1). The analysis revealed that the four most important components (≥5% explained variance; Figure 2B) collectively accounted for ∼70% of the global variance. The first component, Dim.1, explained 37.61% (Figure 2AB) and significantly separated proteome profiles of samples according to the growth phase (Figure 2C). As expected in TSB culture medium, the differences of the proteome profiles in dimension 1 (Dim.1) are predominantly driven by proteins belonging to the CcpA regulon (Figure 2D). CcpA regulates the central metabolism in regard to available glucose and enables the cell to adjust to changing nutrient availability (80). The second component, Dim.2, explained 13.13% of the global variance and separated proteome profiles according to the ClpX status and the iron and oxygen status (Figure 2). Of note, Dim.3 also separated proteome profiles according to the ClpX status as well as the iron and oxygen status (Figure 2C) and explained a similar amount of variance (12.62%). The observed differences were driven by proteins associated with the Rex regulon for Dim.2 and the Fur regulon for Dim.3 (Figure 2D). A weak enrichment of the Rex regulon (GSEA p-value 0.02; Table S2) was also detected for Dim.3. Rex is the master regulator for anaerobic gene expression (81), whereas Fur is the master regulator of the iron limitation response in *S. aureus* (82). Dim.4 was clearly separated by samples solely according to the iron status and expectedly, the differences are driven by the Fur regulon (Figure 2CD).

**Figure 2.**
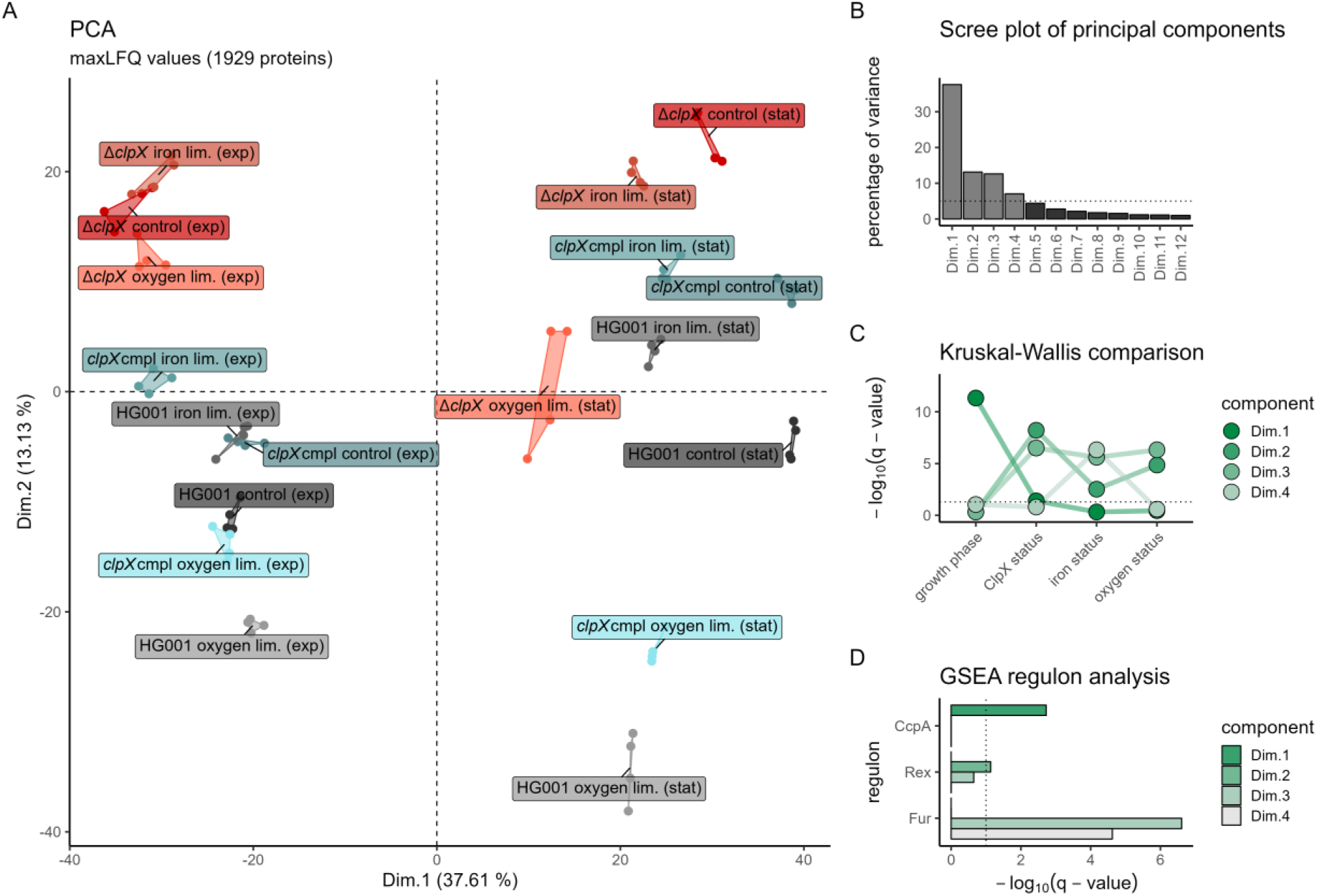
Overview of general proteome profiles. (A) PCA displaying the first and the second component. Each strain and sampling condition was labelled individually. Biological replicates are displayed as points. (B) Scree plot of principal components. The percentage of global variance described by the respective component is displayed for components describing more than 1% of variance. Components describing more than 5% of variance are colored in light grey. The 5% threshold is depicted as dotted line. (C) Separation profiles for the principal components. Negative decadal logarithm of q-values (FDR-adjusted p-values) of Kruskal-Wallis-tests for separation of each condition subcategory (growth phase, ClpX status, iron status, oxygen status) for the most important four principal components are displayed. The q-value threshold for significance of 0.05 is depicted as dotted line. (D) GSEA regulon analysis of proteins spanning the principal components. One-sided GSEA analysis was performed on the percentage of weight of each protein into each principal component for transcription factor regulons according to *Aureo*Wiki. Negative decadal logarithm of q-values (FDR-adjusted p-values) are displayed for the four most important principal components. Regulons with a q-value less than 0.1 (dotted line) for at least one principal component are shown and only regulons with more than five identified members were considered.

The PCA revealed, for the second and third component, an entanglement of the effect of ClpX deficiency and infection-relevant stresses. This finding indicates that a subgroup of proteins specifically synthesized in response to the particular stresses are subject to control by ClpX.

### Identification of a robust ClpX modulon

We aimed to identify the group of proteins directly or indirectly modulated by ClpX irrespective of the stress condition and growth state of the *S. aureus* cell (Figure 3 & Table 2). Hence, we applied strict criteria to obtain a robust ClpX modulon based on the ROPECA statistic (Table S3): (i) protein levels were significantly altered (|fold change| ≥ 1.5 & q-value ≤ 0.05) concordantly between the Δ*clpX* mutant and the HG001 wild-type in both, the exponential and stationary growth phase under the respective condition (iron limitation, oxygen limitation and control); (ii) protein levels were significantly altered concordantly between the Δ*clpX* mutant and the *clpX* complementant in both, the exponential and stationary growth phase under the respective stress condition; (iii) concordant classification as part of the positive or negative ClpX modulon between the three conditions (Table S4A). Using this approach, 24 proteins were defined as members of the robust ClpX modulon (Figure 3A; Table 2). The identified robust ClpX modulon was compared to available data sets (Figure 3B): (i) a global label free proteome analysis comparing *S. aureus* NCTC 8325 and its isogenic Δ*clpX* mutant in stationary phase in rich medium (79) (ii) a transcriptome analysis of *S. aureus* JE2 and its isogenic Δ*clpX* mutant in the exponential growth phase in rich medium (78), and (iii) ClpP trapped proteins in NCTC 8325-4 and Newman in rich medium describing putative direct ClpXP targets (83). Of the 24 ClpX modulon members, 20 were previously identified by at least one of the studies. However, none of them has been identified in all three studies (Figure 3B, Table 2). Of the 24 proteins, 19 proteins were negatively associated with ClpX proficiency (negative ClpX modulon; Figure 3C) and five proteins were positively associated (positive ClpX modulon; Figure 3D).

**Figure 3.**
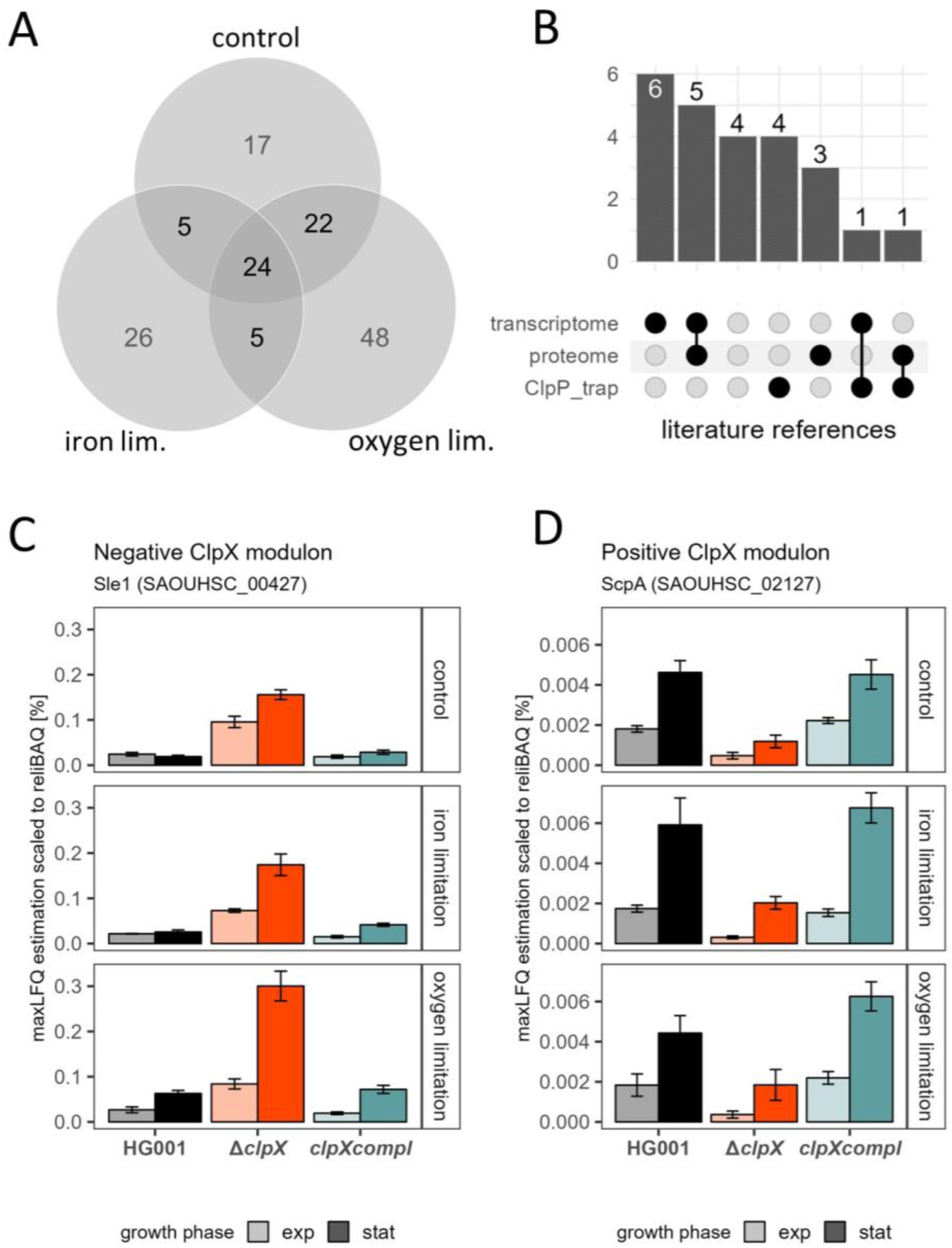
Definition of the robust ClpX modulon. (A) Robust ClpX modulon as overlap of the stress-specific growth phase-independent ClpX modulons. For each condition (control, iron limitation and oxygen limitation), proteins concordantly significantly altered in protein abundance (|fold change| ≥1.5 & ROPECA q-value ≤ 0.05) between the Δ*clpX* mutant and the HG001 wild-type as well as the *clpX* complementant in both the exponential and stationary growth phase were selected. The comparison of the resulting ClpX modulons is represented as a Venn-diagram. (B) Validation of the robust ClpX modulon by literature references. The 24 proteins of the robust ClpX modulon were compared to the previously published ClpX-mediated changes of the transcriptome (78) and of the global proteome (79). In addition, known ClpP-trapped proteins (83) were compared. The intersection of the 24 proteins with the reference studies is represented as an UpSet plot. (C &D) Example of a protein profile belonging to the negative (C) and positive (D) ClpX modulon. Sle1 was detected as a member of the robust ClpX modulon with a negative association with *clpX* proficiency. ScpA was detected as a member of the robust ClpX modulon with a positive association with *clpX* proficiency. The protein level profile is visualized as a barplot depicting the maxLFQ protein level scaled to the relative iBAQ value. No statistics are shown for the barplots as the definition of the robust ClpX modulon was not based on protein level estimation but using the peptide-level ROPECA statistic.

**Table 2.**
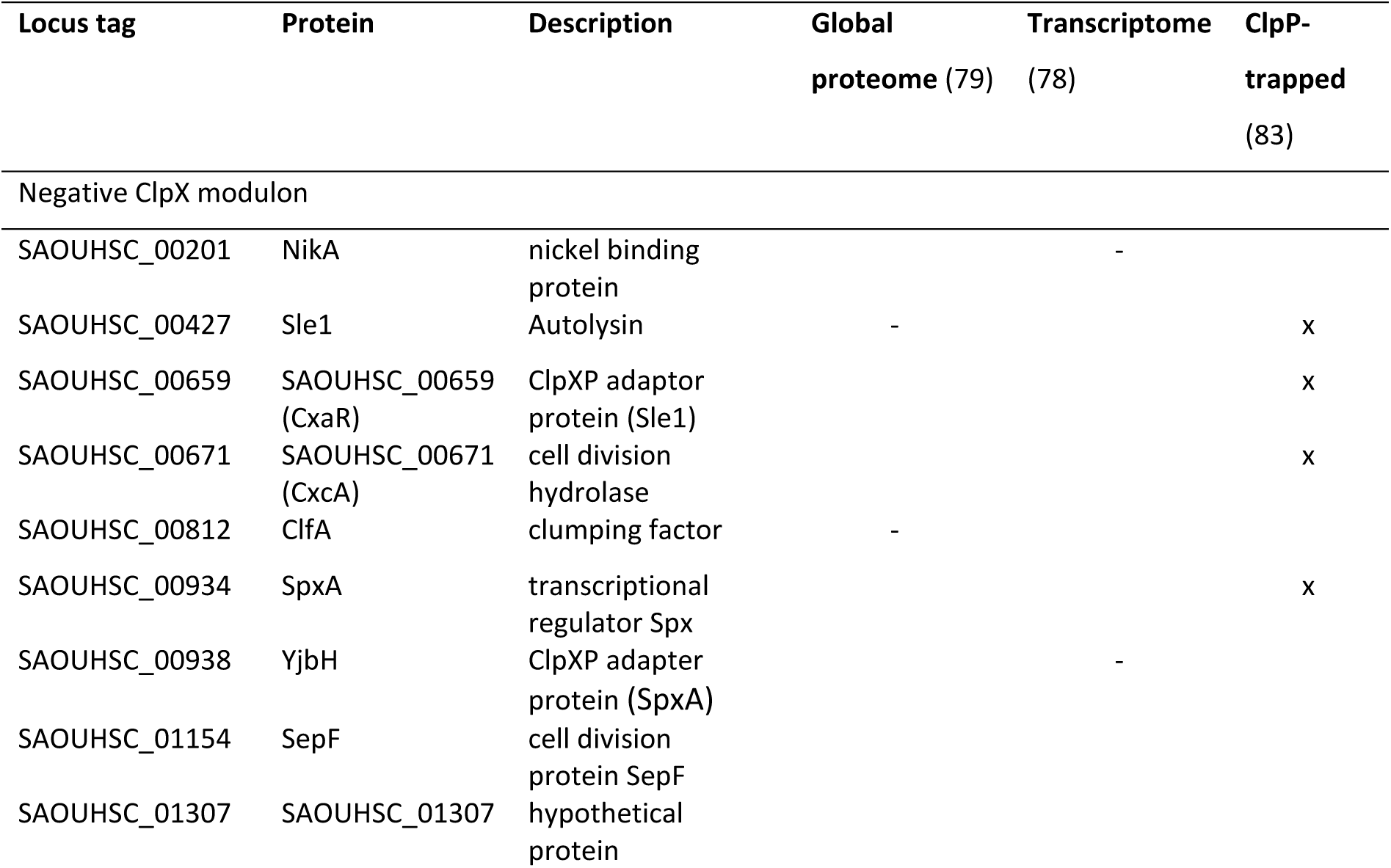
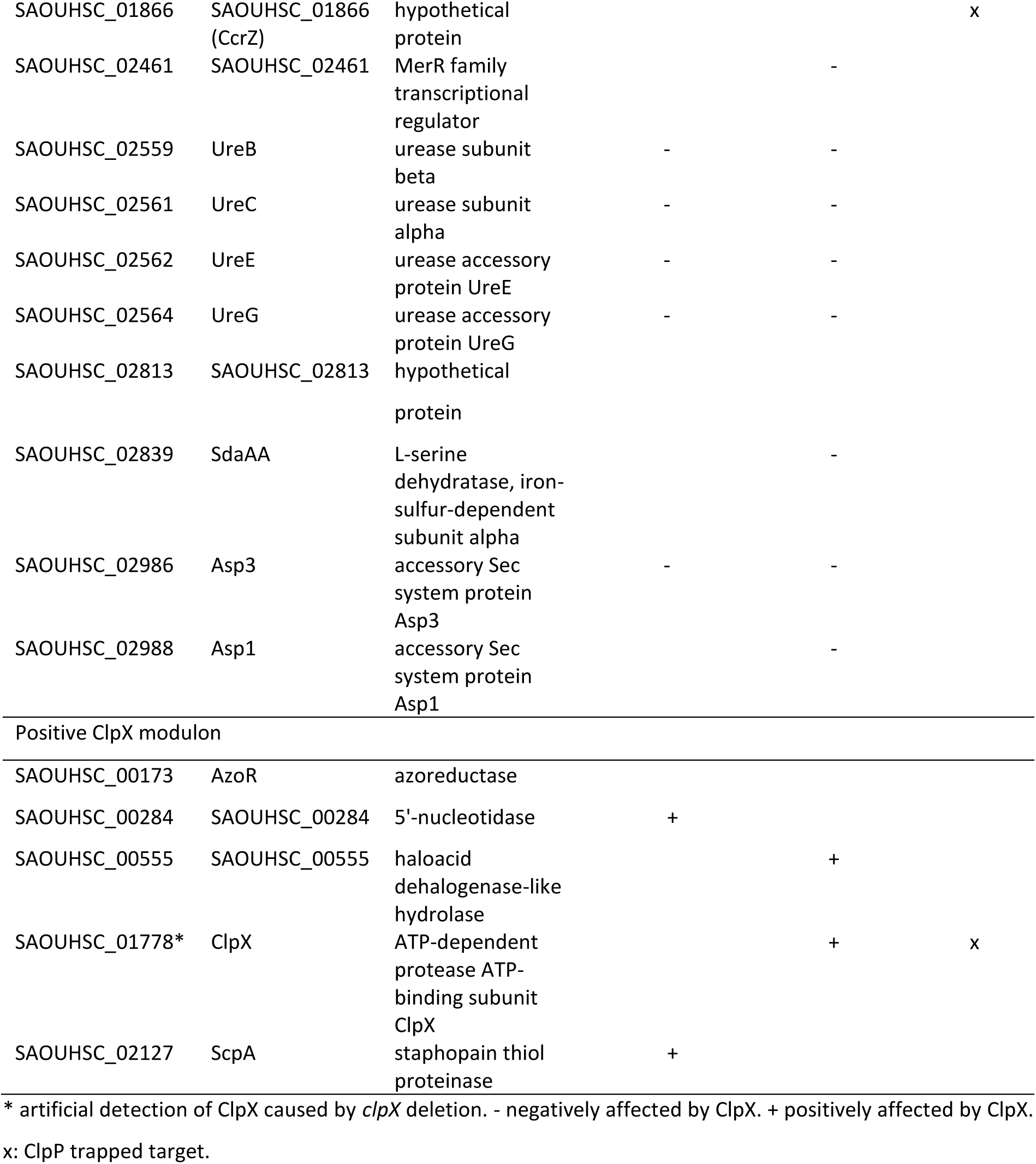
Robust ClpX modulon.

The negative ClpX modulon comprised the well-described target SpxA (12, 84) and its adaptor protein YjbH, which is necessary for ClpXP-mediated proteolysis of SpxA (11). ClpX is also involved in daughter cell splitting, and cell wall metabolism (13, 41, 77, 78), which is partly mediated by the ClpXP-targeted cell wall hydrolases CxcA/SAOUHSC_00671 (14) and Sle1 (Figure 3C; (77, 85, 86)). Both hydrolases and the ClpX adaptor protein SAOUHSC_00659 (CxaR; (87)) were identified as part of the negative ClpX modulon. The cell division protein SepF (88) and the cell cycle protein SAOUHSC_01866 (CcrZ; (89)) were also identified as members of the negative ClpX modulon in line with recent studies showing that ClpX unfoldase activity promotes cell division in *S. aureus* (15, 24, 86). Consistently, the Δ*clpX* mutants compared to the wild-type lead to reduced forward scatter in flow cytometry analysis indicating general reduction of the cell size of ClpX-deficient strains (Figure S20A).

In addition, the urease proteins UreBCEG and the nickel-binding protein NikA were also identified. Urease activity is dependent on nickel and therefore the NikABCDE and NixA nickel transporter act together with the urease in neutralization of low a pH especially during later growth phases (90, 91). The pH of Δ*clpX* cultures at 24 h of cultivation was higher under control and oxygen-limited conditions compared to HG001 cultures (Figure S3).

The positive ClpX modulon member AzoR is one of the two azoreductases of *S. aureus* and both are involved in the quinone stress response (92). However, in contrast to the QsrR-regulation of *azoR1*/SAOUHSC_00320, the regulation of *azoR* is unknown in *S. aureus*. The known QsrR regulon (93), had in general reduced protein levels in the Δ*clpX* mutant (Data S1). In addition, the member of the positive ClpX modulon staphopain A is an extracellular protease degrading several host proteins and shaping the infection process (94, 95).

### Unraveling of independently modulated groups of proteins by an iModulon-like approach to detect general ClpX-mediated effects on the proteome

We demonstrated that the iron and oxygen limitation response and the effect of ClpX deficiency are intermingled in the PCA. Accordingly, we aimed to unravel the intermingling effect of the stress response and the effect of ClpX deficiency. Here, we applied an independent component analysis (ICA) deconvolution method inspired by the iModulon approach originally developed for transcriptomic data sets (70).

We identified 32 so-called i-modulons (Figure 4A; Table S5BC) which explain 90.3% of the observed total variance of the experimental proteomic data set. Explained variance of each single i-modulon (Figure S11B) revealed that the i-modulons 14 (31.5%), 07 (10.3%), 23 (10.2%) and 01 (10.2%) were most important to reflect the proteome profiles, explaining together 62.2% of the total variance. This four i-modulons mirror the applied experimental conditions and demonstrate their importance, as is highlighted in the association of i-modulons with the experimental condition parameters (Figure S11A; Table S5D) and regulatory groups, defined by the *Aureo*Wiki-based regulons, the IsrR (55), the RsaE targetome (96) and the robust ClpX modulon (Figure 4B; Table S5E). I-modulon_14 represents mainly CcpA-driven effects of the growth phase, 07 reflected the mainly Fur-driven effects of iron limitation, 23 represented the mainly Rex-driven effects of oxygen limitation and 01 reflected ClpX-mediated changes of the proteome.

**Figure 4.**
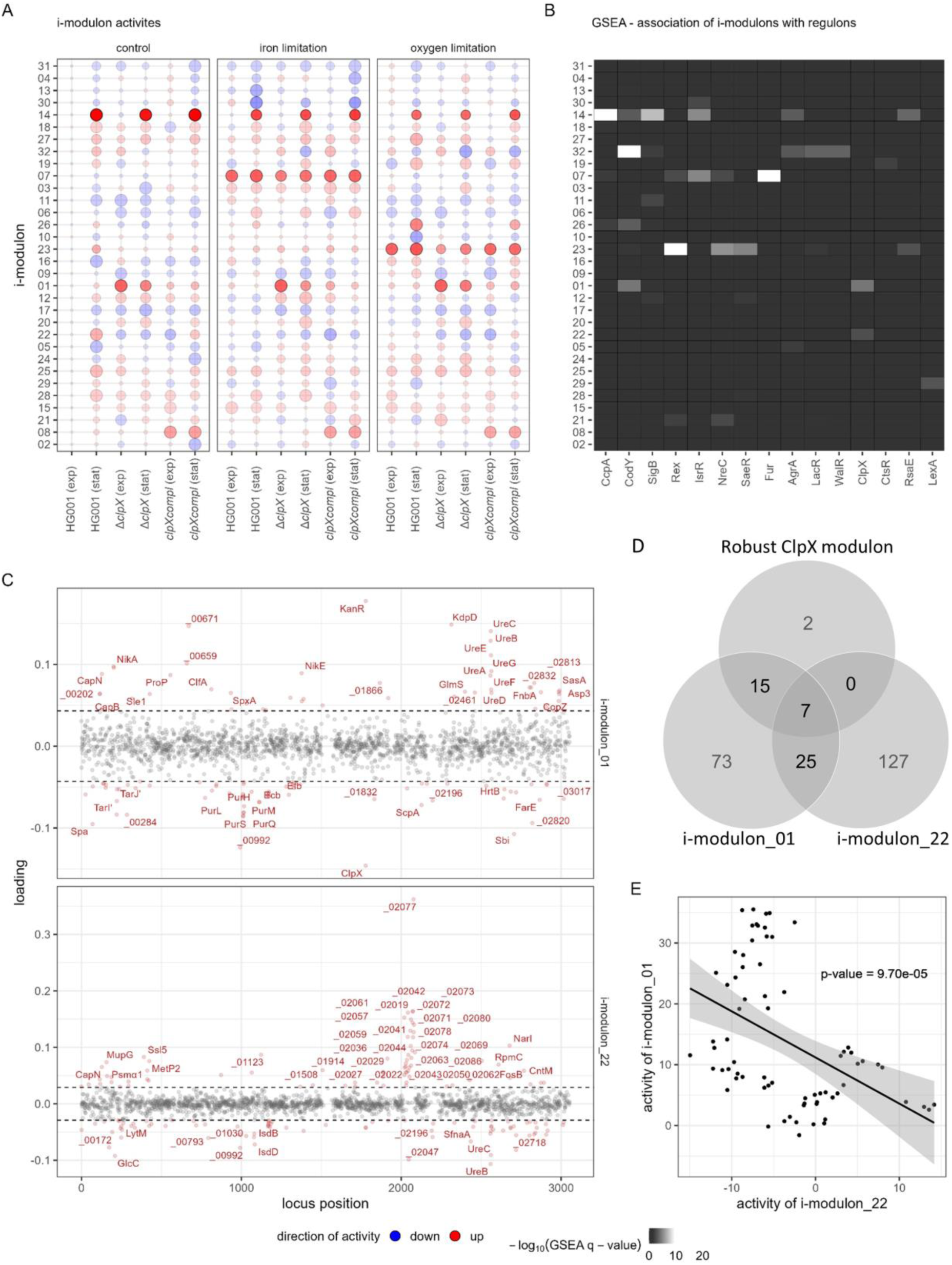
Independent component analysis of the global proteome. (A) Representation of the activity matrix from the ICA. For the four replicates of one condition, the mean activity was calculated. The size of the bubbles represents the activity scaled to the absolute maximum per i-modulon, the transparency indicates the raw activity value and the direction of activity is depicted in color. The detected i-modulons were ordered according to hierarchical clustering (using the 1-|r_pearson_| distance metric; Figure S9). (B) Association of the i-modulons with known regulatory entities. The protein loadings for each i-modulon (matrix M) were tested for enrichment in the high and low weighted proteins using a GSEA. P-values were adjusted using the Benjamini-Hochberg method. Regulon, targetomes or the robust ClpX modulon were considered significantly enriched with q-value ≤ 0.01. (C) Representation of loadings of proteins for the two ClpX associated i-modulons 01 and 22. Protein loadings were plotted to the genome position. Dotted lines indicate loading thresholds for i-modulon members (Figure S10). I-modulon members are indicated in red and the top 50 members were labelled. (D) Venn-diagram representing the overlap of the member sets of the robust ClpX modulon and the two identified ClpX-associated i-modulons. (E) Correlation of i-modulon activities of the two ClpX-associated i-modulons 01 and 22. P-value of Pearson correlation between the activity profiles is indicated.

Twelve i-modulons were associated with the ClpX status (Figure S11A). However, only two of these were also associated with the robust ClpX modulon: i-modulon_01 and i-modulon_22 (Figure 4BC). Therefore, we regarded the protein members of the two i-modulons as potentially ClpX-modulated. Of the 24 robust ClpX modulon members, 22 are members of the i-modulon_01 and seven of these are also members of the i-modulon_22 (Figure 4CD). Examples are the urease subunits UreABCDEFG and the nickel uptake systems components NikADE. The members of i-modulon_01 (Figure 4C) were assessed for overrepresentation of functional annotations (Figure S13B) revealing association with e.g. the urea cycle, purines and pyrimidines as well as virulence. Of note, as ClpX has a negative loading in i-modulon_01, protein loadings are positively associated with ClpX-deficiency and mirror protein levels as in a ClpX-deficient strain. Whereas loadings of the UreABCDEFG proteins were positive, loadings for the i-modulon_01 of the PurCDEFHKLMNQS, PyrBCDEF and CarAB as well as GuaC proteins were negative (Figure 4C, Table S5). In agreement with the model, nucleotide and nucleoside levels are reduced in a ClpX-deficient strain (79). Several virulence factors, such as Spa, Sbi, ScpA, Ecb, Efb, Emp, Eap/Map, ClfA and FnbA, and the regulators SarS and SarV were also identified as i-modulon_01 members (Table S5A). ClpX has been described to interfere with the Agr quorum sensing system activity (19) and expression of several genes encoding the Sar virulence regulator family (27, 28, 97). Protein levels of the Sar virulence regulator family were widely affected by ClpX deficiency (Figure S14, Table S4).

In addition, several proteins, which are part of the cell wall synthesis, are i-modulon_01 members. The CapA, CapB, CapN and CapO proteins, which contribute to the synthesis and anchoring of capsular polysaccharides (98), were positively affected by ClpX deficiency, whereas the alternative wall teichoic acid (WTA) synthesis proteins TarI’, TarJ’ and TarL’ (99) were negatively affected. Intriguingly, inhibitors of WTA synthesis rescue growth of ClpX-deficient cells (13) and lipoteichoic acid (LTA) becomes non-essential in cells lacking ClpX (41). We noted that LtaS and UgtP, which are critical for LTA anchoring (100), were increased in ClpX-deficient cells especially under iron-limited conditions (Table S4, Data S1). In addition, LcpB (SAOUHSC_00997) attaching WTA to the peptidoglycan (101) as well as the N-acetylglucosamine-providing GlmS (102), the peptidoglycan synthetizing proteins MurA2, MurD and MurI (103) and the penicillin-binding proteins Pbp2 and Pbp4 (104) were elevated upon ClpX deficiency (Data 1). Effects of ClpX on peptidoglycan cross-linking and β-lactam resistance has been reported earlier (13, 77, 105).

The second robust ClpX modulon-associated i-modulon, i-modulon_22, showed an overrepresentation of mobile element functions. Indeed, the i-modulon primarily picked up proteins of the prophage φ11 (SAOUHSC_02019 to SAOUHSC_02089; Figure 4C; (106)). Activity of the two ClpX i-modulons was negatively correlated (Figure 4E) and in that term, φ11 activation is negatively associated with ClpX deficiency. The switch between the lysogenic and lytic state of φ11 is controlled by the CI and Cro regulators (107). To induce the lytic state, the N-terminal DNA-binding domain of CI interacts after auto-cleavage with the ClpX chaperone (108), similar to the ClpX-dependent degradation mechanism of LexA (109). I-modulon_22 demonstrates that the ICA unravels, aside general characteristics, complex condition-specific traits of ClpX deficiency as the φ11 activity was especially high in the stationary phase under the control and oxygen-limited condition (Figure 4A).

### ClpX specifically affects the heme uptake system and the staphyloferrin B synthesis and uptake system of the *S. aureus* iron limitation stimulon

The PCA (Figure 3) demonstrated that the effect of the ClpX status and the iron status interplay. As the ICA was suitable to identify condition-specific effects of ClpX deficiency (Figure 4), we inspected the i-modulons associated with the ClpX status and found that i-modulon_12 was overrepresented for iron acquisition proteins (Figure S13B). I-modulon_12 showed a similar activity profile as the ClpX i-modulon_01 (Figure S13A) and investigation of the member proteins revealed negative loadings for IsdA, IsdB, IsdC, IsdD, IsdE and SbnG, indicating a reduction of protein levels upon ClpX deficiency (Figure S13C). ROPECA-based comparisons between the *clpX* mutant and the wild-type as well as the *clpX-*complemented strain revealed that proteins of the heme uptake system (IsdABCDEGHI) and the staphyloferrin B synthesis and uptake system (SbnABCEFGHI and SirA) were decreased in the mutant under iron limitation (Figure 5A & S15B, Table S3 and S4).

**Figure 5.**
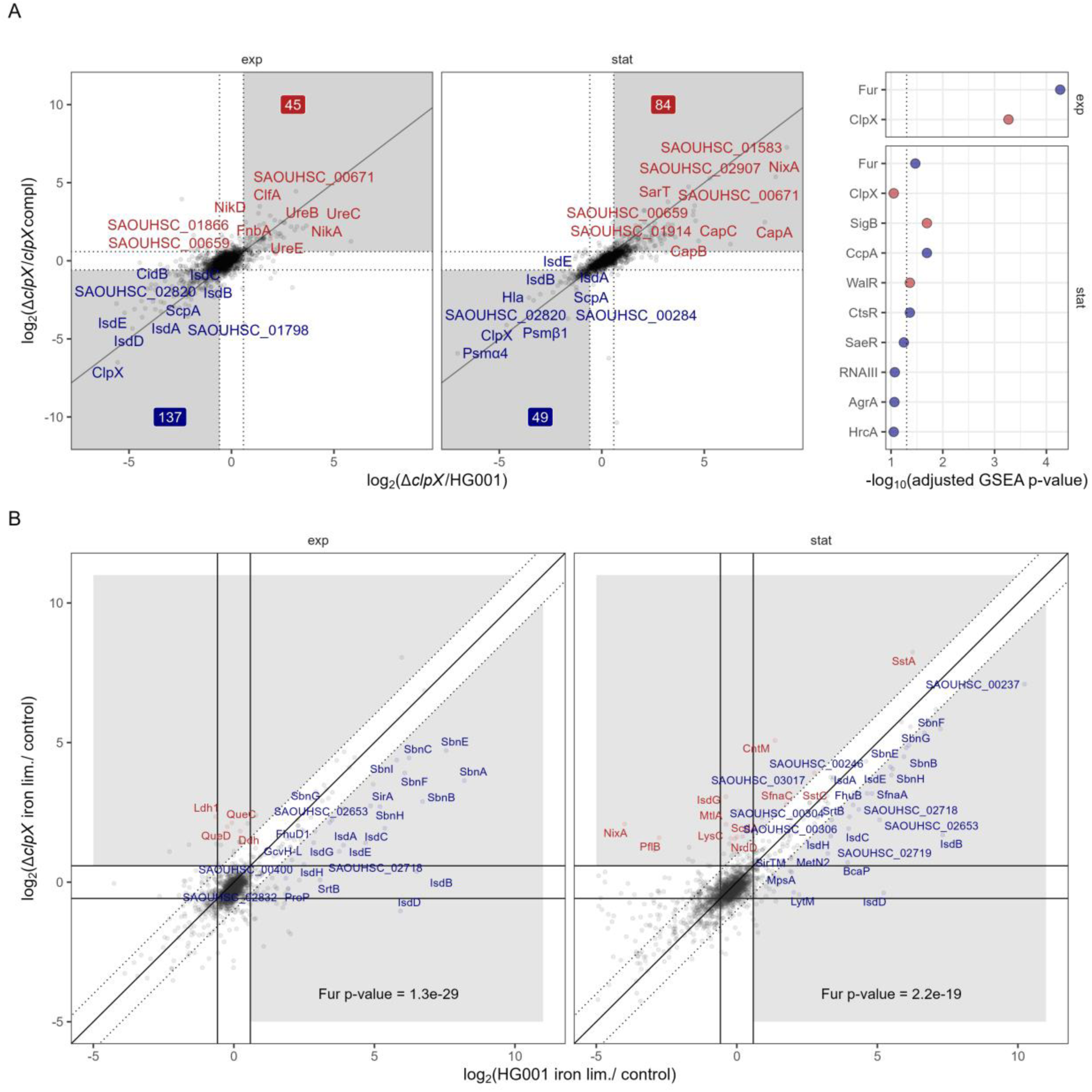
ClpX affects induction of parts of the Fur regulon under iron-limited conditions. (A) ClpX modulated proteins under iron limitation. ROPECA pairwise statistics were used. Proteins altered in the Δ*clpX* mutant compared to the HG001 wild-type and the complemented strain were visualized. Top ten proteins, which were significantly altered (|fold change| ≥ 1.5 & q-value ≤ 0.05) in both comparisons were labelled. The fold change thresholds are indicated as dotted lines. Significantly reduced protein levels are indicated in blue and significantly elevated protein levels are indicated in red. Numbers indicate the total number of significantly reduced and elevated proteins. Comparisons for the exponential and stationary growth phase are separated. GSEA statistics for the analysis are visualized in the right panel. Positive enrichment scores are indicated in red and negative scores are indicated in blue. The dotted line indicates the 0.05 threshold. Proteins were ranked according to the signed Euclidean distance to the zero point considering the log_2_ ratios and q-values. (B) Proteins induced by iron limitation in the wild-type and the Δ*clpX* mutant. Perfect correlation of induction is indicated as solid line and 2-fold deviations are indicated as dotted lines. Proteins with 2-fold deviation and significant induction (fold change≥ 1.5 & q-value ≤ 0.05) in at least one of the strains were labelled. Proteins with higher induction in the wild-type are labelled red and proteins with higher induction in the mutant are labelled blue. Overrepresentation of the Fur regulon was tested using the Fisher’s Exact test.

In line with this, the Clp system has a broad impact on the iron limitation response (26, 79). More specifically, for *clpX* and *clpP* mutants, Farrand *et al*. observed missing induction of *isdB*, encoding the hemoglobin receptor IsdB, under iron limitation and resulting reduced heme acquisition capacities (25). We detected an increase in the heme uptake system as well as the staphyloferrin B synthesis and uptake system proteins under iron limitation compared to control conditions in both the wild-type and the Δ*clpX* mutant with a lesser extent in the Δ*clpX* mutant (Figure 5B). The effect of ClpX deficiency on induction of the heme uptake system was more pronounced than on the staphyloferrin B synthesis and uptake system, which is in agreement with the fact that the staphyloferrin B system and siderophore utilization are regulated by available heme levels (110–112). As we recently reported a similar phenotype upon inactivation of sRNA IsrR (55), we tested if the IsrR levels are affected in the *clpX* mutant, however, no effect was observed (Figure S16A). In addition, Fur and the Fur antagonist protein Fpa (113) did not display altered levels in the ClpX deficient-strain either (Figure S16BC), which is consistent with the Fur-independent regulation of the *isdB* transcript level in the *clpX* mutant (25). Expression of the *isd* genes and the genes involved in the staphyloferrin B uptake are also induced by different oxidative stresses (114–116). The thiol- and quinone-stress regulators QsrR (93), HypR (115) and SpxA showed altered protein levels in the Δ*clpX* mutant (Figure S17). We also observed lower protein level induction of the *sirTM* operon encoding a sirtuin/macrodomain system (SAOUHSC_00304, GcvH-L, SAOUHSC_00306, SirTM, LplA2; Figure 5B), which is likely involved in the intracellular survival and oxidative stress response (117–119).

### ClpX interferes with the energy metabolism especially under oxygen limitation

In a similar manner as for the iron limitation response, the PCA revealed an intermingling effect of the oxygen limitation response and the effect of ClpX deficiency in the second and third component (Figure 2). Thus, we asked how ClpX interacts with the oxygen limitation response. Intriguingly, the i-modulon_23, which is the main i-modulon reflecting the oxygen limitation response, showed less activity for the ClpX-deficient strain compared to the ClpX-proficient strains (Figure 4A).

In agreement, the *clpX* mutant did show reduced induction of the Rex regulon upon oxygen limitation compared to the wild-type (Figure 6AB & S15C). Rex is the master regulator of the oxygen limitation stimulon (81). Since no ClpX-dependent effect on Rex protein levels were observed (Data S1), the difference in the Rex regulon activity indicated differences in the NAD^+^/NADH ratio of the *clpX* mutant compared to the ClpX-proficient strains especially under oxygen limited conditions (81). In the Gram-positive bacterium *Corynebacterium crenatum*, *clpX* deletion positively affected the NADP^+^/NADPH ratio during fermentation (120). In *S. aureus*, an influence of ClpX on fermentation or the oxygen limitation response has not been described yet. However, for a Δ*clpP* mutant reduction of Rex subregulon was shown (20) and importantly, ClpP but not the second unfoldase ClpC is essential for fermentative growth (23), suggesting ClpX to play an important role in fermentation.

**Figure 6.**
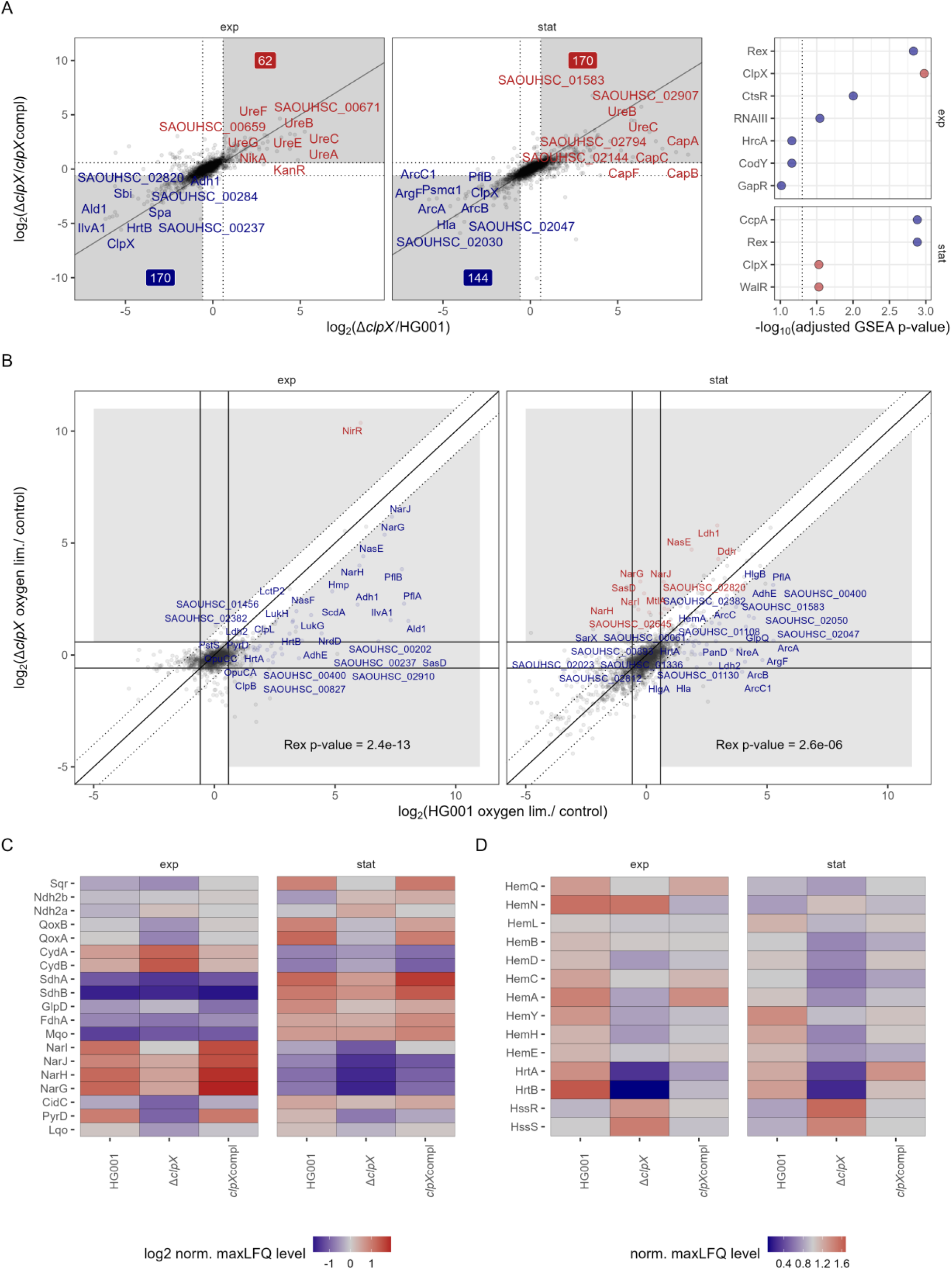
ClpX affects the energy metabolism under oxygen limitation. (A) ClpX modulated proteins under oxygen limitation. ROPECA pairwise statistics were used. Proteins altered in the Δ*clpX* mutant compared to the HG001 wild-type and the complemented strain were visualized. Top ten proteins, which were significantly altered (|fold change| ≥ 1.5 & q-value ≤ 0.05) in both comparisons were labelled. The fold change thresholds are indicated as dotted lines. Significantly reduced protein levels are indicated in blue and significantly elevated protein levels are indicated in red. Numbers indicate the total number of significantly reduced and elevated proteins. Comparisons for the exponential and stationary growth phase are separated. GSEA statistics for the analysis are visualized in the right panel. Positive enrichment scores are indicated in red and negative scores are indicated in blue. The dotted line indicates the 0.05 threshold. Proteins were ranked according to the signed Euclidean distance to the zero point considering the log_2_ ratios and q-values. (B) Proteins induced by oxygen limitation in the wild-type and the Δ*clpX* mutant. Perfect correlation of induction is indicated as solid line and 2-fold deviations are indicated as dotted lines. Proteins with 2-fold deviation and significant induction (fold change ≥ 1.5 & q-value ≤ 0.05) in at least one of the strains were labelled. Proteins with higher induction in the wild-type are labelled red and proteins with higher induction in the mutant are labelled blue. Overrepresentation of the Rex regulon was tested using the Fisher’s Exact test. (C) Heat map of proteins involved in the respiratory chain. Mean maxLFQ values were centered to the median and log2 transformed. (D) Heat map of proteins involved in the heme homeostasis. Mean maxLFQ values were centered to the median.

Of note, the two component system SrrAB, which is also involved in the oxygen limitation response in *S. aureus*, is a Rex regulon member (81) and accordingly showed reduced levels in the ClpX-deficient strain compared to the proficient strains (Data S1). SrrAB is in particular involved in the regulation of the respiratory chain and in particular the *qoxABCD*, *cydAB*, and *hemACDX* genes (121).

The likely differences in the NAD^+^/NADH ratio also implicated differences in the activity of the respiratory chain. *S. aureus* contains three terminal electron acceptor reductases, namely CydBA, QoxABC (oxygen respiration; (122)) and NarGHIJ (nitrate respiration; (123)), as well as ten quinone reductases (FdhA-complex, SdhCAB, Sqr, GlpD, CidC, PyrD, Mqo, Lqo, Ndh2a, Ndh2b (124)). We observed broad effects of ClpX deficiency on the respiratory chain proteins (Figure 6C). There seems to be a connection with a recently described link between the molybdopterin biosynthesis and ClpXP (125) promoting nitrate respiration and FdhA activity (126, 127). However, no accumulation of the suggested ClpXP-target MoeA in the ClpX-deficient strain was observed. In addition, the respiratory chain is also affected by heme levels as SdhC (128), QoxABCD, CydAB (122) and NarI (129) use heme as a cofactor. We noted that proteins of the heme biosynthesis were reduced in the *clpX* mutant and that the effect was especially pronounced for HemA (Figure 6D), which is critical for regulation of heme biosynthesis (130, 131). Even though Clp-dependent regulation of the heme biosynthesis has not been described in *S. aureus*, there are examples of such regulation in other organisms (132–134).

We also noted that the heme efflux subunit HrtB (135) was negatively influenced by the ClpX deficiency according to i-modulon_01 (Figure 4C). Indeed, HrtA and HrtB protein levels were drastically reduced in the *clpX* mutant (Figure 6D) indicating low heme stress in the mutant compared to both the wild-type and the complemented strain under oxygen limitation. *HrtAB* expression is under control of the two component system HssRS sensing the intracellular heme level and is expressed under heme toxicity conditions (136). Interestingly, HssRS showed higher protein levels in the *clpX* mutant compared to the wild-type and complemented strain (Figure 6D). In the Gram-positive bacterium *Bacillus anthracis*, ClpX has been reported to be required for full HssRS activity (137).

### ClpX determines the virulence of *S. aureus* in *Galleria mellonella* and supports intracellular replication of *S. aureus* in human bronchial epithelial cells

As demonstrated by the proteomic profiles under infection-relevant conditions, a plethora of proteins, which are associated with virulence and bacterial fitness, are dysregulated with regard to their abundance in the ClpX-deficient strain. In a systemic *G. mellonella* infection model (Figure 7A &S19), the Δ*clpX* mutant showed a significant attenuation compared to the wild-type, as determined by the mortality number of *Galleria* larvae over a period of ten days post-infection (p.i.). Larvae infected with the Δ*clpX* mutant revealed a significantly higher survival rate than those infected with the HG001 wild-type (p-value log-rank test: 0.014). Virulence of the Δ*clpX* mutant was reduced to approximately half of that of the wild-type (HR Δ*clpX* vs. HG001: 0.47). The Δ*clpX* mutant nevertheless exhibited a certain degree of virulence compared to the 0.9% NaCl control (p-value log-rank test: 0.002; HR Δ*clpX* vs. 0.9% NaCl control: 3.99).

**Figure 7.**
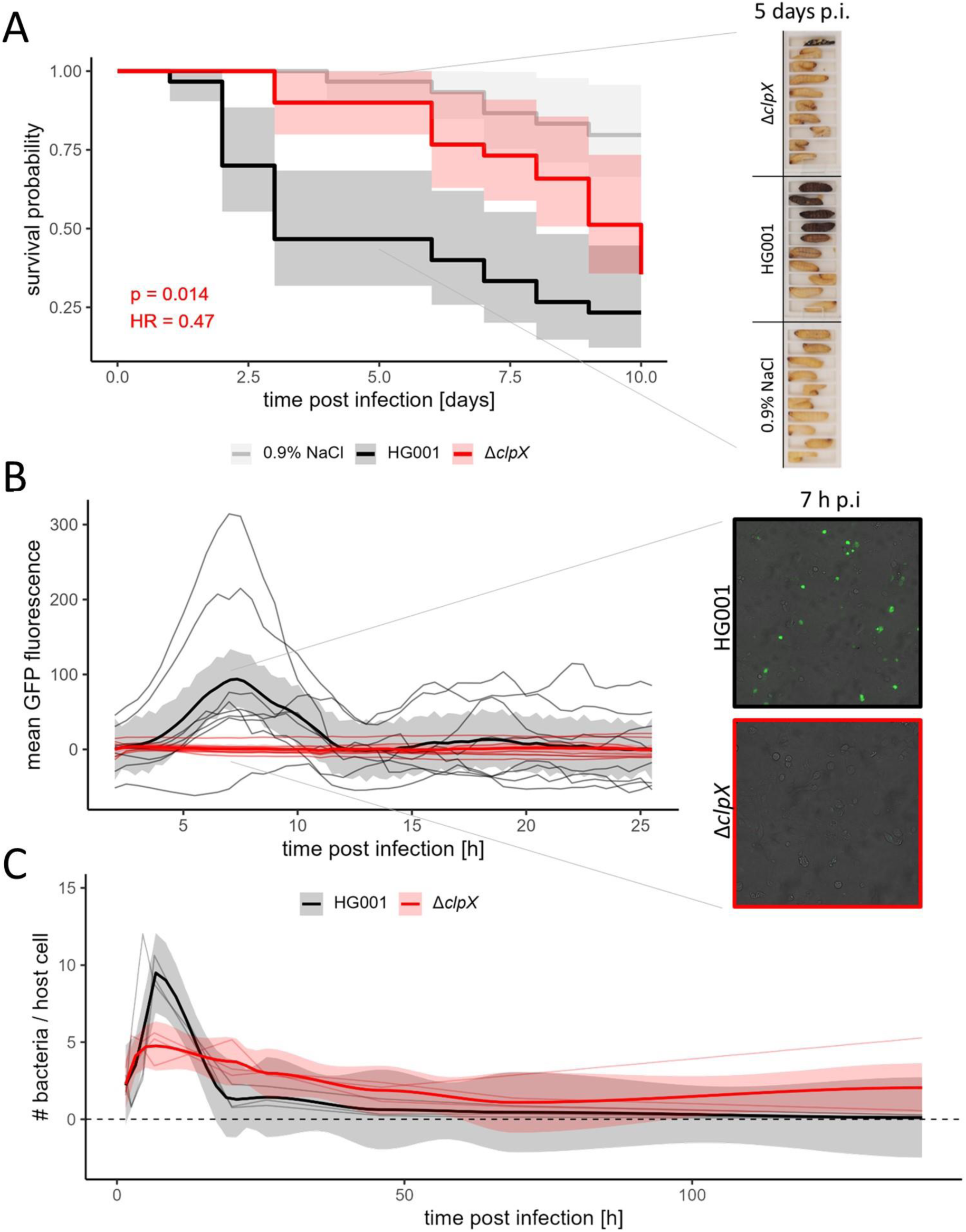
ClpX is an essential factor for *S. aureus* pathogenicity. (A) *Galleria mellonella* infection experiment. *G. mellonella* larvae were infected with 1×10^5^ *S. aureus* cells in 10 µl 0.9% NaCl solution or as treatment control 10 µl sterile 0.9% NaCl solution was injected. Ten larvae per bacterial strain (HG001 wild-type and Δ*clpX* mutant) or treatment control and biological independent experiments were infected. In total, in three biological replicates 30 larvae per strain were infected. Upper track: Survival was fitted to Kaplan-Meier survival estimates and difference between survival of larvae infected with the Δ*clpX* mutant and survival of larvae infected with the HG001 wild-type was tested using a log-rank test (p-value indicated). 95% confidence intervals (CI) of the Kaplan-Meier fit are depicted. The hazard ratio (HR) comparing Δ*clpX* vs. HG001 infection was estimated using a Cox proportional hazards regression model. Lower track: Representative larvae of one independent biological replicate five days post infection (p.i.). (B) Quantification of intracellular *S. aureus* cells based on GFP fluorescence intensity during live cell imaging of infected in 16HBE14o-cells (MOI = 40). GFP was tracked from 2 h p.i. to 26 h p.i. every 30 min. Mean GFP signal per picture was quantified for two biological replicates with each three technical replicates. GFP signal time curves were MOI-normalized and background corrected and the loess fit of all replicates per bacterial strain was depicted in bold with the corresponding 95% CI. Zoom: Representative microscopy pictures 7 h p.i. (C) Quantification of intracellular *S. aureus* cells *via* flow cytometry-based cell counting in 16HBE14o-cells (MOI = 50). Intracellular bacteria were counted at 1.5 h, 2.5 h, 4.5 h, 6.5 h, 20 h, 26 h, 46 h, 69 h and 140 h p.i. The number of bacterial cells were MOI-normalized and normalized to the number of host cells. Three biological replicates were obtained. Loess fit of the replicates per bacterial strain is depicted in bold with the corresponding 95% CI.

*S. aureus* possesses the specific trait of being an extracellular as well as an intracellular pathogen (138). Intracellular infection of *S. aureus* in relation to ClpX-deficiency was investigated using 16HBE14o-bronchial epithelial cells (Figure 7BC & 21). In accordance with Kim *et al.* (30), we did not observe significant differences in internalization between the ClpX-proficient and -deficient strains (Figure S20B). Live cell imaging tracked from 2 h post infection (p.i.) up to 26 h p.i. (Figure 7B), demonstrated a rapid increase of the fluorescence signal up to 7 h post infection for the HG001 wild-typereflecting the quantity of intracellular GFP-labelled bacterial cells. Rapid multiplication of the HG001 wild-type in non-professional phagocytic host cells during the early phase of infection is in line with previous studies (17, 48). In contrast, for the Δ*clpX* mutant, drastically reduced intracellular multiplication was tracked during this early phases of infection (Figure 7B &S21). Similar, ClpX-driven effects on intracellular replication during short-term infections were previously reported (24, 30). However, when following the intracellular behavior of the two strains for up to 140h p.i. by cell counting and microscopy (Figure 7C & Figure S21), we observed that during later stages of the infection (after 24h p.i.) the number of bacteria per host cells was higher for the Δ*clpX* mutant compared to the wild-type HG001. This could indicate that a high proportion of the ClpX-deficient strain enters an intracellularly persisting state, in which the bacteria survive but no multiplication occurs.

## Discussion

We demonstrated that ClpX plays a decisive role in *S. aureus* pathophysiology and is critical for protein homeostasis of a plethora of infection-relevant physiological aspects (Figure 8).

**Figure 8.**
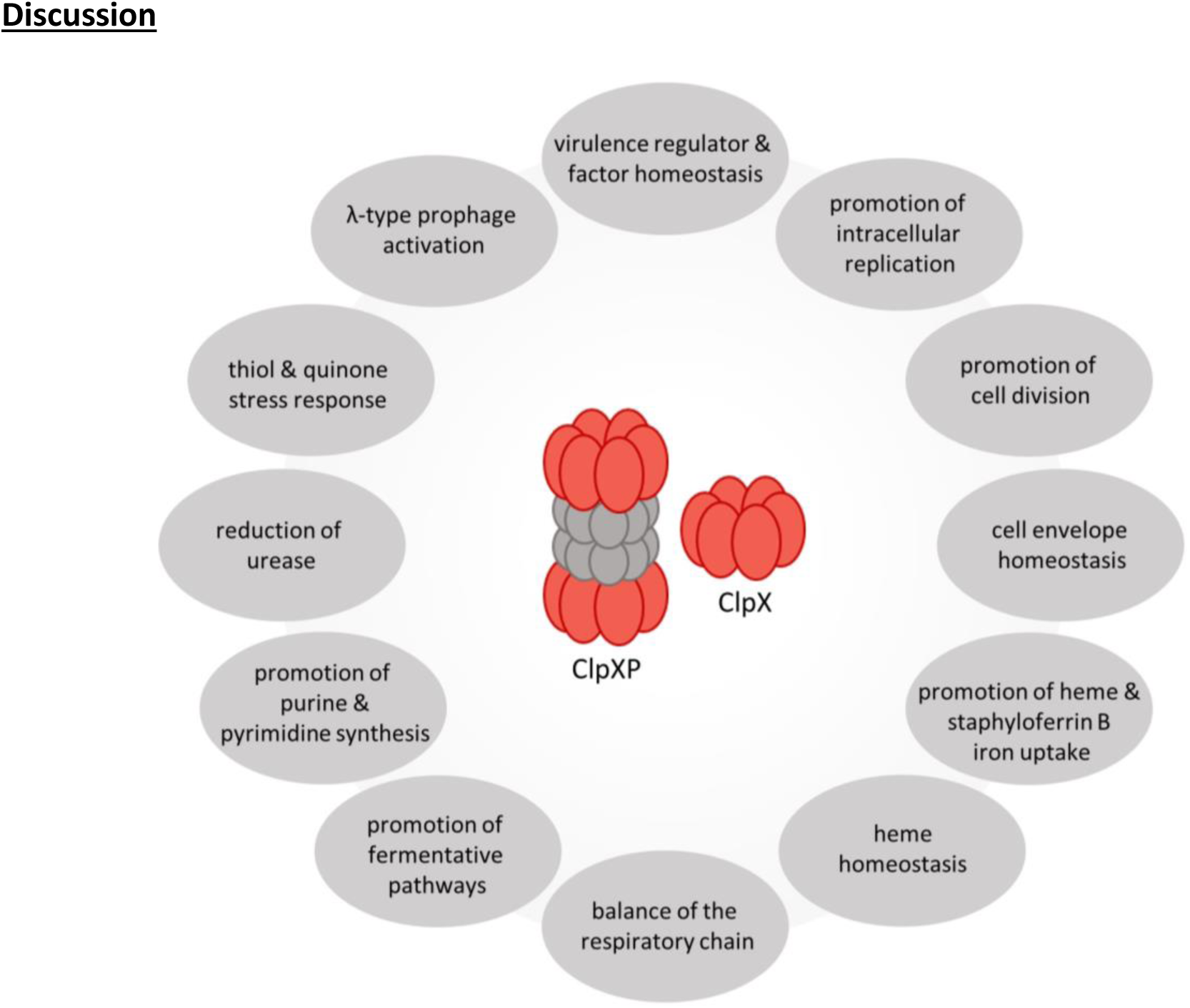
Summary of ClpX-driven effects on *S. aureus* physiology and pathophysiology.

In the systemic *G. mellonella* model, ClpX deficiency led to strongly reduced virulence and comparable observations were made in mice models with *clpX* and *clpP* deletion mutants (19, 25, 29, 30, 85, 139). In skin and pneumonia infection models, ClpX-deficient strains caused reduced host immune responses (29, 30). Intriguingly, in our cell culture infection experiments, the ClpX-deficient strains showed reduced intracellular replication during early stage of infection (24, 30) and a high proportion of intracellularly persisting bacterial cells. Interestingly, *clpC* deletion also enhances bacterial intracellular persistence (140). The high proportion of persisting cells raises the question, if reduced ClpX levels support persister formation and recurring infection processes. Previously, a decline of ClpX protein levels has been observed in long-term intracellular *S. aureus* (17), supporting the strategy of persister formation during infection.

Additionally, *clpX* is known as a intrahost evolution hot-spot (141, 142) and one mechanism of host immune evasion is the acquisition of *clpP* mutations (143, 144).

Persistence can occur passively by inhibited escape of *S. aureus* cells from the phagosomes or actively by down-regulation of the metabolism and cell division in the cytoplasm (145). Several virulence factors such as PSMα, PSMβ, δ-toxin and β-toxin are associated with the escape of internalized *S. aureus* from the phagosome (146, 147). Accordingly, the regulators SarA and Agr are required for phagosomal escape (148–150). Our data has reflected the known role of ClpX role in decreasing the Agr and Sar family system activity (27). Strikingly, ScpA, which is part of the robust positive ClpX modulon, induces host cell death and reduced levels of ScpA lead to prolonged intracellular survival of *S. aureus* (149, 151).

However, several other proteins, which are not considered to be classical virulence factors were also dysregulated in the *clpX* mutant and were previously associated with cytotoxicity (150, 152). This highlights the link between bacterial fitness, virulence and intracellular persistence. Among these, SecA2, TarL’, PurB, PurF and SbnF were also dysregulated in the *clpX* mutant. SecA2 and the two negative robust ClpX modulon members Asp1 and Asp3 are part of the accessory Sec system of *S. aureus* (153, 154). Interestingly, the signal peptidase I, SpsB, was also increased in protein levels in the *clpX* mutant, which is in line with the observed dysregulation of several cell envelope associated proteins and virulence factors in our study (155). Additionally, retention and release of secreted proteins is in part also mediated by PG, LTA and WTA (156, 157), whose synthesis and homeostasis are also affected by ClpX deficiency.

Long-term survival of *S. aureus* in the host is clearly also driven by the metabolism (48, 158). For example, *de novo* purine and pyrimidine biosynthesis is critical for bacterial replication in infection models (159, 160) and the *ure* operon is expressed as part of the weak acid stress response and is critical for persistence during infections (91). Reduction of the purine and pyrimidine synthesis and increase of the urease activity are hallmarks of the Δ*clpX* phenotype.

*S. aureus* cells persisting in infections are associated with the small colony variant (SCV) phenotype which is defined by slowed growth (161). As described, ClpX deficiency led to slightly reduced growth rates even in TSB and the cell division proteins SepF and CcrZ were identified as negative ClpX modulon members. In line with the SCV and persister phenotype, ClpX inactivation has been associated with higher antibiotic resistance (77, 105, 162, 163). The prolonged survival of SCVs is supported by a low membrane potential, which is, e.g., the result of a reduced TCA or an impaired respiratory chain (161). Interestingly, ClpXP was recently identified to be involved in lipid homeostasis (162) and proteins of the respiratory chain were dysregulated in the ClpX-deficient strain. However, in contrast to most stable SCVs (164), the Δ*clpX* mutant showed a reduced induction of the anaerobic metabolism in our data. Further, mutants in several *hem* genes cause non-respiring SCVs (164). Intriguingly, proteins involved in heme synthesis and efflux systems were reduced in the Δ*clpX* mutant.

Under iron limitation, the heme uptake system was less induced in the Δ*clpX* mutant. This uptake system is especially shaping the infection processes of *S. aureus* (25, 165). Interestingly, inactivation of MspA controlling LTA biosynthesis (166) was reported to lead to elevated levels of both, the heme uptake system and the heme efflux system in *S. aureus* (167). This demonstrates that the control of LTA homeostasis by ClpX might also be critical for the heme homeostasis.

In this study, we provide a global state-of-the-art proteomic data set as a resource allowing further insights into the role of ClpX deficiency in *S. aureus* pathophysiology. We also demonstrated that the precise consequences of Clp system deficiency for the infection process and the development of persistence still need to be unraveled. Our study highlighted that ClpX is involved in the iron and oxygen limitation response as central aspects of bacterial fitness in infection processes. As ClpXP is one of the most conserved protease systems in pathogenic bacteria (5, 6), the here provided information likely can be also transferred to other pathogenic organisms. The Clp system is often considered as potential targets for therapeutic purposes (e.g. 168–170). However, the potential role of Clp deficiency in persisting infections might lead to question its suitability as a drug target.

## Supporting information

Combined Supplemental Data

## Availability of data and materials

MS data is deposited to the MassIVE repository under accession MSV000095845 (ProteomeXchange dataset PXD055808).

## Acknowledgments

We thank Marc Schaffer, Emilia Schmidt and Gina Wockenfuß for excellent assistance with the experiments, Anna Nagel for the evaluation of the influence of codon optimization of fluorochromes, Leif Steil for providing the heavy-labelled library and the support with MassIVE. Language was revised using the DeepL Write (www.deepl.com/de/write) tool.

