## Supplementary material for "Unraveling proteomic chaos by independent component analysis - ClpX proficiency promotes the iron and oxygen limitation responses of *Staphylococcus aureus* and affects the intracellular bacterial behavior": Combined Supplemental Data

### Supplemental Data Index

### 1 Supplementary Methods

#### Method A. Construction of the empty allele exchange plasmid pSauSE.

For the construction of the backbone of the pSauSE allele exchange plasmid, the plasmid pBASE6 (Geiger *et al.*, 2012) was optimized regarding the vector size and the utilization of selection markers. Primers used for the cloning approaches are listed in Supplemental Data Table A.

pSauSE was constructed using the SLIC method (Li and Elledge, 2012) in *E. coli* Stellar$^{TM}$ (TaKaRa Bio, Japan) cells. The temperature-sensitive *S. aureus* pE194$_{ts}$ origin of replication (Luchansky and Faith, 1991; Brückner, 1997) and the *E. coli* pBR322 origin of replication (Bolivar *et al.*, 1977; Brückner, 1997), as well as the AHT-inducible *secY*-asRNA system as plasmid counter selection system (Bae and Schneewind, 2006) were exploited from the pBASE6 plasmid. The promoter p$_{help}$ (Riedel *et al.*, 2007; Monk *et al.*, 2012) was cloned upstream of the *cat* gene from pBASE6 allowing the use of this selection marker in *E. coli* and *S. aureus*. In addition, an *Eco*RV restriction site (Kholmina *et al.*, 1980) was integrated allowing for linearization of the empty plasmid. The EcoRV (Thermo Fisher Scientific, USA)-digested pSauSE plasmid was used for construction of specific allele exchange plasmids. A schema of the plasmid is shown in Figure S1B.

#### Method B. Construction of the plasmid pJL-sar-GFP$_{redopt}$.

The GFP-labeling plasmid pJL-sar-GFP$_{redopt}$ was constructed in *E. coli* Stellar$^{TM}$ cells using the SLIC method (Li and Elledge, 2012). The Gfpmut2-encoding sequence from the plasmid pJL74 (Liese *et al.*, 2013) was replaced with an *S. aureus* codon-optimized variant of *gfpmut2.* Here, codon usage was optimized to the tRNA sequence number distribution in *S. aureus*. The codon-optimized *gfp* variant was synthetized (GenScript, USA). Primers used for the cloning approaches are listed in Supplemental Data Table A. A non-codon optimized *gfp* variant was shown to induce the CodY-regulated amino acid starvation response by usage of low frequent amino-acyl-tRNA species (Figure S18). The codon-optimized variant pJL-sar-GFP$_{redopt}$ did not induce the amino acid starvation response compared to the non-plasmid-carrying and an empty control plasmid-carrying strain.

#### Method C. Eukaryotic cell cultivation.

The human epithelial cell line 16HBE14o-, a transformed bronchial epithelial cell line (Cozens *et al.*, 1994) obtained from Prof. J.-P. Hildebrandt (University of Greifswald, Greifswald; (Möller *et al.*, 2020)), was employed for infection experiments. Until infection, the cells were maintained at 37 °C in a humidified environment with 5% $CO_2$ in eukaryotic minimal essential medium (eMEM): 1x MEM (Biochrom AG, Germany) supplemented with 10% (v/v) fetal calf serum (FCS; Biochrom AG, Germany), 2% (v/v) 200 mM L -glutamine (PAN-Biotech GmbH, Germany) and 1% (v/v) 100x nonessential amino acids (PAN-Biotech GmbH, Germany). Cells were passaged every three to four days using trypsin-EDTA for detachment of the cells (PAN-Biotech GmbH, Germany).

#### Method D. *Galleria mellonella* growth conditions and infection model.

*Galleria mellonella* larvae (proinsects GmbH, Germany) were incubated for one to two days in the dark at room temperature and sorted regarding the weight and fitness upon arrival. Insects were fed using a mixture of oat bran (125 g), oat flakes (30 g), skimmed milk powder (5 g), dried yeast (5 g), honey (25 g) and glycerol (as needed). The method for the infection experiment was adjusted from (Tsai, Loh and Proft, 2016; Trevijano-Contador and Zaragoza, 2018; Wojda *et al.*, 2020; Ménard, Rouillon, Cattoir, *et al.*, 2021).

*G. mellonella* larvae, the larvae of the greater wax moth (Wojda *et al.*, 2020), are recently used as a time- and cost-efficient model host to study host-pathogen interactions in a wide range of pathogens (Piatek, Sheehan and Kavanagh, 2021). Results are comparable to results determined in mammalian infection models (Jander, Rahme and Ausubel, 2000; Brennan *et al.*, 2002; Sheehan, Dixon and Kavanagh, 2019) as the insect immune system is similar to the mammalian innate immune system (Sheehan *et al.*, 2018) and therefor the *Galleria* infection model can serve as a preliminary model to understandstaphylococcal infection processes (Sheehan, Dixon and Kavanagh, 2019; Ménard, Rouillon, Ghukasyan, *et al.*, 2021; Alnezary *et al.*, 2023).

#### Method E. $^{15}$N external standard-based normalization of peptide ion intensities.

To produce the in-house heavy labelled $^{15}$N protein standard as described in (Busch *et al.*, 2025; Illenseher *et al.*, 2025), *B. subtilis* BSB1 cells (Nicolas *et al.*, 2012) were precultivated in a 1:10 dilution series up to the 10$^{th}$ dilution in BioExpress® Bacterial Cell Media (U-$^{15}$N, 98%; Cambridge Isotope Laboratories Inc., USA) aerobically overnight at 37°C. The tube with the highest dilution level with visible growth was used to inoculate the main culture in the heavy labelling medium to an initial $OD_{540nm}$ of 0.05. The culture was harvested at mid-exponential growth phase ($OD_{540nm}$ = 1.5) by centrifugation (3 min at 4°C and 8,500 xg) and the *B. subtilis* cell pellets were washed with 20 mM HEPES pH 8.0. The pellets were lysed as described in the methods of the main manuscript using a bead mill, followed by SDS denaturation of the proteins at 95°C and subsequent Benzonase digestion to eliminate nucleic acids. Protein concentration of the heavy labelled standard was also determined as described in the methods of the main manuscript using the Micro BCA™ Protein Assay Kit.

To ensure reliable quantification of protein levels, we normalized the peptide ion intensity based on the external standard (Figure S6) instead of cross sample normalization methods such as the median-median normalization because we noted broad differences in the peptide ion composition (Figure S5). The different experimental conditions induced altered protein compositions and profiles under the infection-relevant stress, the different ClpX status and growth states which then lead to the differences in peptide ion composition. Of note, we could generally observe higher levels of explained total ion currents for stationary growth phase samples (Figure S6) and higher numbers of identified peptide ions in the Δ*clpX* mutant in the stationary growth phase (Figure S5), which is in line with a slight general protein accumulation in this strain (Stahlhut *et al.*, 2017; Alqarzaee *et al.*, 2021). Of note, however, ClpC likely has a more pronounced function than ClpX in the general proteolysis of *S. aureus* (Chatterjee *et al.*, 2011; Stahlhut *et al.*, 2017), which is in line with the observed slight increase.

We also asked if ClpX deficiency might lead to an increased proportion of oxidized proteins either by protein stress or by the lack of protein quality control resulting is accumulation of nonfunctional proteins (Ezraty *et al.*, 2017; Lévy *et al.*, 2019; Schramm, Schroeder and Jonas, 2020; Matavacas and von Wachenfeldt, 2022). In contrast to previous findings (Alqarzaee *et al.*, 2021), the proportion of methionine oxidized peptide ions did not change in intensity nor in identification numbers (Figure S7). Possibly, such stress responses occur in later phases of cell survival.

#### Method F. Background on the iModulon-like approach.

The iModulon approach has been introduced by (Sastry *et al.*, 2019; Rychel *et al.*, 2021) and has been applied successfully on several transcriptomic data sets of *e.g*., *E. coli* (Anand *et al.*, 2019; Sastry *et al.*, 2019; Tan *et al.*, 2020; Lamoureux *et al.*, 2023; Rychel *et al.*, 2023), *B. subtilis* (Rychel, Sastry and Palsson, 2020; Sastry *et al.*, 2024), *S. aureus* (Poudel *et al.*, 2020, 2022; Fait *et al.*, 2022; Gao *et al.*, 2023; Yu *et al.*, 2023), *Pseudomonas aeruginosa* (Rajput *et al.*, 2022) to identify biological meaningful independently modulated sets of genes reflecting modulons or in best case regulons (Qiu *et al.*, 2024). This approach has been demonstrated to be robust in clustering of similar genes across several complex data sets (Sastry *et al.*, 2021) and in principle transferable to proteomic data sets (Patel *et al.*, 2024).

Classical hierarchical clustering approaches of the protein patterns only allow for assigning proteins to one regulator mechanism because each protein belongs to only one cluster (Figure S8). However, analyses of this type do not completely fulfill the need for unraveling a complex biological regulatory network, where multiple regulators or regulator entities can act on one gene or protein. In contrast, as i-modulon activities are considered to be linear and additive, the iModulon approach can assign one protein to several i-modulons which is reflecting the complexity of biological networks.

The main idea of ICA is to identify statistical independent components in a data set and thereby disentangle mixed signals. In case of the presented data set, the rel_iBAQ-rescaled maxLFQ protein intensities were considered to be a mixed signal ($X \in M^{\#proteins \times \#samples}$) of several modulons with different activities in the samples. The modulons are represented by identified independent i-modulons and their i-modulon members (proteins). One protein might not necessarily be a member of only one i-modulon because protein intensity can also biologically result from multiple additive modulon activities.

In the ICA, it is assumed that (i) an activity signal $A \in M^{\#i\text{-}modulons \times \#samples}$ (i-modulon activity per sample) is solely mixed by $M \in M^{\#proteins \times \#modulons}$ (loadings of proteins comprising to the i-modulons), (ii) that the effect of regulation between the i-modulons is statistical independent, (iii) that the i-modulons are non-Gaussian distributed, and (iv) that the effects of different regulations are linearly additive. This leads to the model: X = M × A. One is then interested in finding an estimation of M and A, respectively M\` and A\`, so that the statistical independence of the i-modulons in A\` is maximized by maximization of the non-Gaussian distribution of M\` with $M^{`-1} \times X = A`$. Of note, the estimations A\` and M\` cannot predict the scaling of the i-modulon activity and loadings of proteins per modulon and therefore, activities and loadings cannot be directly compared across i-modulons but only within one i-modulon.

Of the 32 i-modulons, 25 were significantly associated with at least one of the applied condition parameters (Figure S11A; Kruskal-Wallis q-value ≤ 0.05) and 11 were significantly associates with at least one of previously known regulatory groups (Figure 4B; GSEA q-value ≤ 0.01). This resulted in total in 27 assigned i-modulons (Table S5F). Inspection of the assigned i-modulons revealed, that rather than regulons, stimulons and modulons were separated (Figure 4B & Figure S12BC).

The ICA aims to identify statistically independent non-Gaussian distributed components, to identify i-modulon members, proteins comprising to non-Gaussian distribution of each i-modulon were selected. Inspection of the i-modulons in regard to enriched known regulons revealed, that rather than perfect single regulons, stimulons and modulons were picked up (Figure 5D & Figure S10BC). We therefor did not define i-modulon groups based on F1-statistcs of known regulatory entities (Sastry *et al.*, 2019) but used the Bonferroni-adjusted p-value of the D'Agostino omnibus test as indicator for reached Gaussian distribution by exclusion of i-modulon members from the independent component (Figure S9). 967 proteins were assigned to at least one i-modulon and each i-modulon group had on average 180.1 members (min: 59, max: 322). No relation between the explained variance and the size of the i-modulon could be observed. The i-modulons reduced to their identified members could still explain 71.4% of the total variance.

As previously stated (Patel *et al.*, 2024), the ICA approach picked up proteins with a high CV (Figure S10A), which means that proteins differing between conditions are grouped into i-modulons. However, the stable proteins could be considered as an additional group, which we labeled i-modulon_00 (Table S5A). Overrepresentation analysis demonstrated that the members widely reflect the association of the whole i-modulon with known regulatory groups (Figure S10B).

### 2 Supplementary Tables

#### Table A. Oligonucleotides used for mutant and plasmid constructions.

| **pSauSE plasmid backbone construction** | | |
|---|---|---|
| **N°** | **Name** | **Sequence** |
| 1 | pSauSE_FragA_for | ATGCTTGATATCTAGAGCGTAAGCGAAAGTAG |
| 2 | pSauSE_FragA_rev | AAAGCATAATGGGATCCCACTTTATCCAATTTTCGTTTG |
| 3 | pSauSE_FragB_for | TAGACAATTGGAAGAGAAAAG |
| 4 | pSauSE_FragB_rev | TCTAGATATCAAGCATTGGTAACTGTCAGAC |
| 5 | pSauSE_FragC_for | ATCCCATTATGCTTTGGCAG |
| 6 | pSauSE_FragC_rev | CTCTTCCAATTGTCTAAATC |
| **SLB001 HG001 *clpX*::*kanR* mutant generation & pSauSE_Δ*clpX*::*kanR* plasmid construction** | | |
| 7 | Ins.Ctrl.DclpX_for | AACTGTTACTGTTCCTGCAG |
| 8 | Ins.Ctrl.clpX_rev | CACCAATAATGATAGCTTGC |
| 9 | pSauSE_linear_for | ATCTAGAGCGTAAGCGAAAGTAG |
| 10 | pSauSE_linear_rev | ATCAAGCATTGGTAACTGTCAG |
| 11 | DclpX_pSauSE_up_for | TTACCAATGCTTGATCAACGCTTTGGTGTAGAAGC |
| 12 | DclpX_Sau_up_rev | GTCATTTTAAACTCCTCCTCAATTCGTTAAAGACTATACTAGATTGG |
| 13 | DclpX_Sau_Km_for | TGAGGAGGAGTTTAAAATGACTAAAATG |
| 14 | Km_just_rev | TTAGAAGAGTTCATCGAGTAAGATG |
| 15 | Km_screen_rev | CCAGCCATAGCATCATGTCC |
| 16 | clpX_Sau_do_for | CTTACTCGATGAACTCTTCTAACATCAGCTTAATCATTGATGTG |
| 17 | clpX_pSauSE_do_rev | TTCGCTTACGCTCTAGATGACCGCAGCATAAGACACAC |
| **pTripleTREP_clpX plasmid construction** | | |
| 18 | pTripleTREP_linear_for | TTCGTTCCACTGAGCGTCAG |
| 19 | pTripleTREP_clpX_rev_long | CTTTTTACACCCCTATTTCCCTATCACTGATAGGGAATCTATC |
| 20 | clpX_TS_SD_for | AATAGGGGTGTAAAAAGAATGTTT |
| 21 | clpX_Term_pTripleTREP_rev | GCTCAGTGGAACGAAAAAAAAGCTCCGATCAAAGTTAAAC |
| 22 | clpX_screen_pTripleTREP_rev | CAACATCATCGCCTACATAAC |
| **pJL-sar-GFP$_{redopt}$ plasmid construction** | | |
| 23 | pJL_linearize_for | GGCGCGCCTATTCTAATGC |
| 24 | pJL_linearize_rev | AAATAATCATCCTCCTAAGAATTC |
| 25 | GFP_NEW_4_pJL74_for | TAGGAGGATGATTATTTATGAGCAAAGGTGAAG |
| 26 | GFP_NEW_4_pJL74_rev | TAGAATAGGCGCGCCTTATTTATACAACTCATCCATACC |

#### Table B. Reversed phase liquid chromatography (RPLC).

| ***Instrument*** | **Ultimate 3000 RSLC (Thermo Scientific)** |
|---|---|
| *Trap column* | 75 µm inner diameter, packed with 3 µm C18 particles (Acclaim PepMap100, Thermo Scientific) |
| *Analytical column* | Accucore 150-C18, (Thermo Fisher Scientific)<br>25 cm x 75 µm, 2.6 µm C18 particles, 150 Å pore size |

| | |
|---|---|
| *Buffer system* | binary buffer system consisting of 0.1% acetic acid in HPLC-grade water (solvent A) and 100% ACN in 0.1% acetic acid (solvent B) |
| *Flow rate* | 300 nl/min |
| *Gradient* | 0 min: 2% B<br>2 min: 5% B<br>10 min: 7% B<br>70 min: 25% B<br>75 min: 40% B<br>77 min: 90% B<br>83 min: 90% B<br>85 min: 2% B<br>95 min: 2% B |
| *Column oven temperature* | 40 °C |

Table C. Mass spectrometry.

| ***Instrument*** | ***Orbitrap Exploris 480*** |
|---|---|
| *Electrospray* | Nanospray Flex$^{TM}$ Ion Source |
| *Operation mode* | data-independent |
| ***Full Scan Properties*** | |
| *MS scan resolution* | 120,000 |
| *AGC target* | 3e6 (300%) |
| *maximum ion injection time for the MS scan* | 60 ms |
| *Scan range* | 350 to 1,200 m/z |
| *Microscans* | 1 |
| *Polarity* | Positive |
| *RF Lens* | 50% |
| *Spectra data type* | Profile |

| *DIA Properties (MS2)* | |
|---|---|
| *Resolution* | 30,000 |
| *maximum ion injection time for the MS/MS scans* | Auto |
| *Normalized AGC target* | 3E6 |
| *Spectra data type* | Profile |
| *Microscans* | 1 |
| *Number of isolation window* | 66 |
| *Isolation window width* | 13 m/z |
| *Window overlay* | 2 m/z |
| *Fixed first mass* | 200 |
| *HCD collision energy* | 30% |

Table D. Spectronaut$^{TM}$ parameters used for data analysis of mass spectrometry data.

| ***Parameter*** | **Setting** |
|---|---|
| ***Calibration*** | |
| *Calibration Mode* | Automatic |
| *RT Regression Type* | Local (Non-Linear) Regression |
| ***Identification*** | |
| *Pvalue Estimator* | Kernel Density Estimator |
| *Precursor Qvalue Cutoff* | 0.001 |
| *Protein Qvalue Cutoff* | 0.01 |
| *Decoy method* | Mutated |
| *Decoy Limit Strategy* | Dynamic |
| ***Library Filters*** | |
| *Fragment Ins* | |

| | |
|---|---|
| *Ion AA Length* | N=3 |
| *m/z* | Min: 200 Max: 3000 |
| *Precursors* | |
| *Best N Fragments per Peptide* | Min: 3 Max: 6 |
| ***Protein Inference*** | |
| *Inference Algorithm* | IDPicker |
| ***Quantification*** | |
| *Precursor Filtering* | Identified (Qvalue) |
| *Imputation Strategy* | Use Background Signal |
| *Quantity MS Level* | MS2 |
| *Quantity Type* | Area |
| *Cross-Run Normalization* | True |
| *Normalization Filter Type* | Library Name Filter |
| *Library Name* | 20230811_Bacillus_heavy_lib_fix_mod |
| *Normalization Strategy* | Global Normalization (Median) |
| *Row Selection* | Identified in all Runs (Complete) |
| *Interference Correction* | True |
| *Only Identified Peptides* | True |
| *Exclude All Multi-Channel Interferences* | True |
| *MS1 Min* | 2 |
| *MS2 Min* | 3 |
| ***Workflow*** | |
| *Multi-Channel Workflow Definition* | From Library Annotation |
| *Profiling Strategy* | iRT Profiling |

| | |
|---|---|
| *Profiling Target Selection*<br>*Unifiy Peptide Peaks Strategy* | Profile only non-identified Precursors<br>Select corresponding Peak |
| ***XIC Extraction*** | |
| *XIC Extraction Window* | Dynamic |
| *MS1 & MS2 Mass Tolerance Strategy* | Dynamic |
| ***Library set up 2024_HG001_DclpX_InfMimicking*** | |
| *Software version* | 18.6.231227.556995 |
| *Digest Rule* | Trypsin/P |
| *Digest Type* | Specific |
| *Missed Cleavage* | 2 |
| *Min Peptide Length* | 7 |
| *Max Peptide Length* | 52 |
| *Toggle N-terminal M* | True |
| *Protein & Peptide FDR* | 0.01 |
| *Fragment Ions per Peptide* | 6-10 |
| *Protein Database* | Saureus_NCTC8325_aug2023_aureowiki_Lysostaphin_Benzonase_Trypsin_Cm_TetR_RsbU_Km.fasta |
| ***Library set up 20230811_Bacillus_heavy_lib_fix_mod*** | |
| *Software version* | 18.1.230626.50606 |
| *Digest Rule* | Trypsin/P |
| *Digest Type* | Specific |
| *Missed Cleavage* | 2 |

| | |
|---|---|
| *Min Peptide Length* | 7 |
| *Max Peptide Length* | 52 |
| *Toggle N-terminal M* | True |
| *Protein & Peptide FDR* | 0.01 |
| *Fragment Ions* | 6-10 |
| *Protein Database* | 2021_01__uniprot_sp_Bsubtilis_168_incl_isofor |
| *Protein Database* | ms.fasta |

**Table E.** R packages for analysis and visualization of the proteome data.

| Package | Version | Reference |
|---|---|---|
| Tidyverse | 2.0.0 | (Wickham *et al.*, 2019) |
| FactoMineR | 2.4 | (Lê, Josse and Husson, 2008) |
| Ggpubr | 0.6.0 | Alboukadel Kassambara (2023). ggpubr: 'ggplot2' Based Publication Ready Plots |
| Ggrepel | 0.9.5 | Kamil Slowikowski (2021). ggrepel: Automatically Position Non-Overlapping Text Labels with 'ggplot2'. |
| Ggtext | 0.1.2 | Claus O. Wilke and Brenton M. Wiernik (2022). ggtext: Improved Text Rendering Support for 'ggplot2'. |
| Iq | 1.9.6 | (Pham, Henneman and Jimenez, 2020) |
| Openxlsx | 4.2.5 | Philipp Schauberger and Alexander Walker (2023). openxlsx: Read, Write and Edit xlsx Files. |
| Patchwork | 1.2.0 | Thomas Lin Pedersen (2024). patchwork: The Composer of Plots. |
| PECA | 1.30.0 | Tomi Suomi, Jukka Hiissa and Laura L. Elo (2021). PECA: Probe-level Expression Change Averaging. |
| Readr | 2.1.4 | Hadley Wickham, Jim Hester and Jennifer Bryan (2023). readr: Read Rectangular Text Data. |
| Readxl | 1.4.3 | Hadley Wickham and Jennifer Bryan (2023). readxl: Read Excel Files. |
| Rstatix | 0.7.2 | Alboukadel Kassambara (2023). rstatix: Pipe-Friendly Framework for Basic Statistical Tests. |
| RColorBrewer | 1.1-3 | Erich Neuwirth (2022). RColorBrewer: ColorBrewer Palettes. |
| Gghalves | 0.1.4 | Frederik Tiedemann (2022). gghalves: Compose Half-Half Plots Using Your Favourite Geoms. |
| Vroom | 1.6.5 | Jim Hester, Hadley Wickham and Jennifer Bryan (2023). vroom: Read and Write Rectangular Text Data Quickly. |
| ComplexUpset | 1.3.3 | (Krassowski, Michal *et al.*, 2022) |

### 3 Supplementary Figures

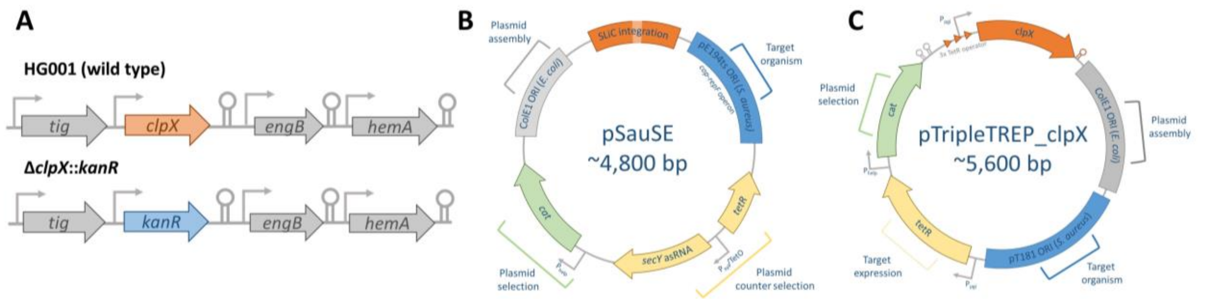

**Figure S1. Construction of the allele exchange strain HG001 Δ*clpX*::*kanR* and the *clpX* complemented strain.** (A) Schematic representation of the chromosomal *clpX* region in the strains HG001 and the allele exchanged HG001 Δ*clpX::kanR*. (B) Schematic representation of the empty mutagenesis plasmid pSauSE. (C) Schematic representation of the complementation plasmid pTripleTREP_clpX.

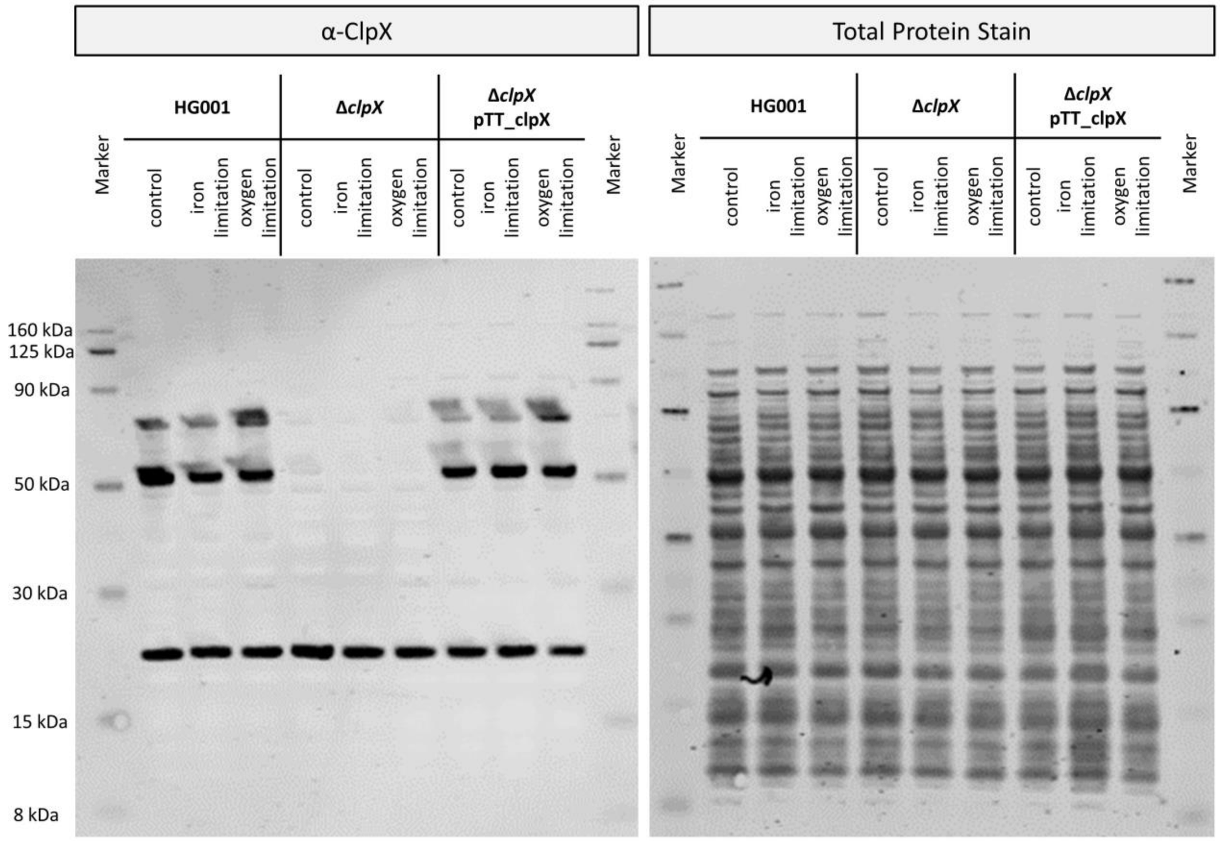

**Figure S2. Western Blot of ClpX.** Representative samples for the three strains under infection-relevant conditions tested are shown. Samples were harvested in exponential growth phase (2.5 h). 5 µg of total protein were used. Left panel: Complete membrane of the

Western Blot shown in Figure 1B using the α-ClpX$_{B. subtilis}$ primary antiserum. Right panel: Complete membrane of the Western Blot shown in Figure 1B using LI-COR Revert Total Protein Stain as loading control.

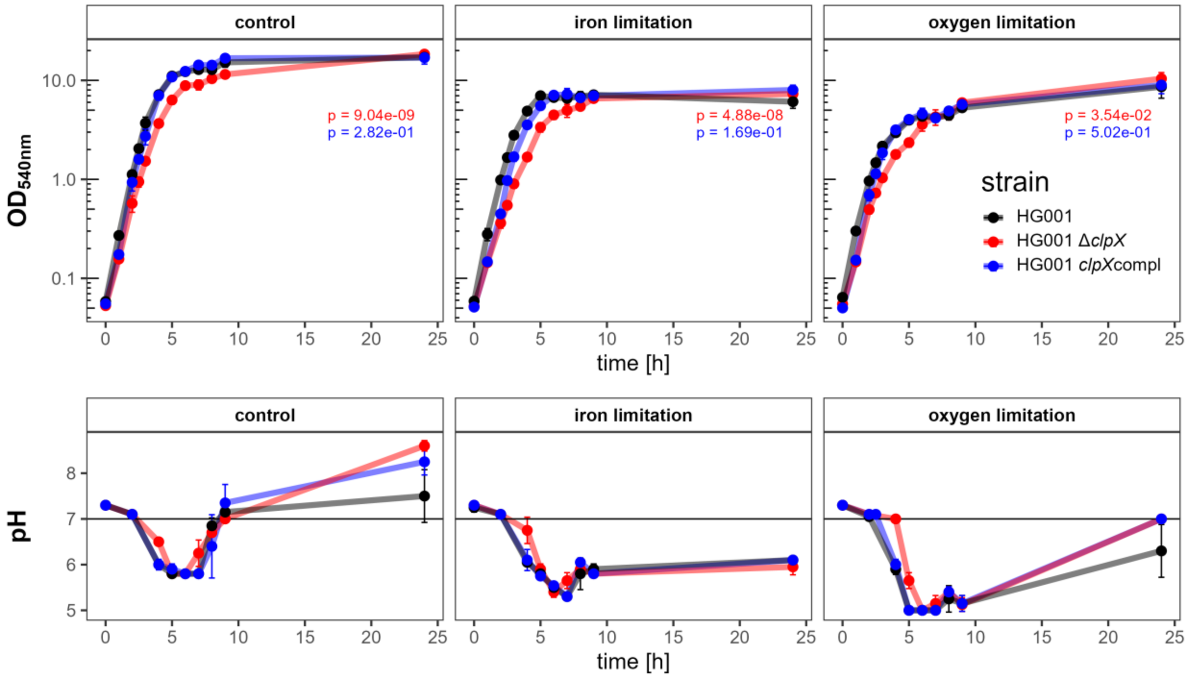

**Figure S3. Cultivation of HG001, Δ*clpX* and *clpX*compl under infection-relevant conditions.** Upper lane: Growth curves. The strains HG001, Δ*clpX* and *clpX*compl were cultivated under control condition (TSB), iron limitation (TSB + 600 µM DP) and oxygen limitation (TSB 100% flask volume). All media contained 20 ng/ml AHT. Growth was hourly measured by $OD_{540}$. Four independent biological replicates were cultivated, standard deviation is indicated as error bars. Differences in general growth was tested by time point-paired t-test between the HG001 wild type and Δ*clpX* (p-value depicted in red) and between the HG001 wild type and *clpX*compl (p-value depicted in blue) per cultivation condition. Lower lane: pH curves. The pH was measured at selected time points. Four independent biological replicates were cultivated, standard deviation is indicated as error bars.

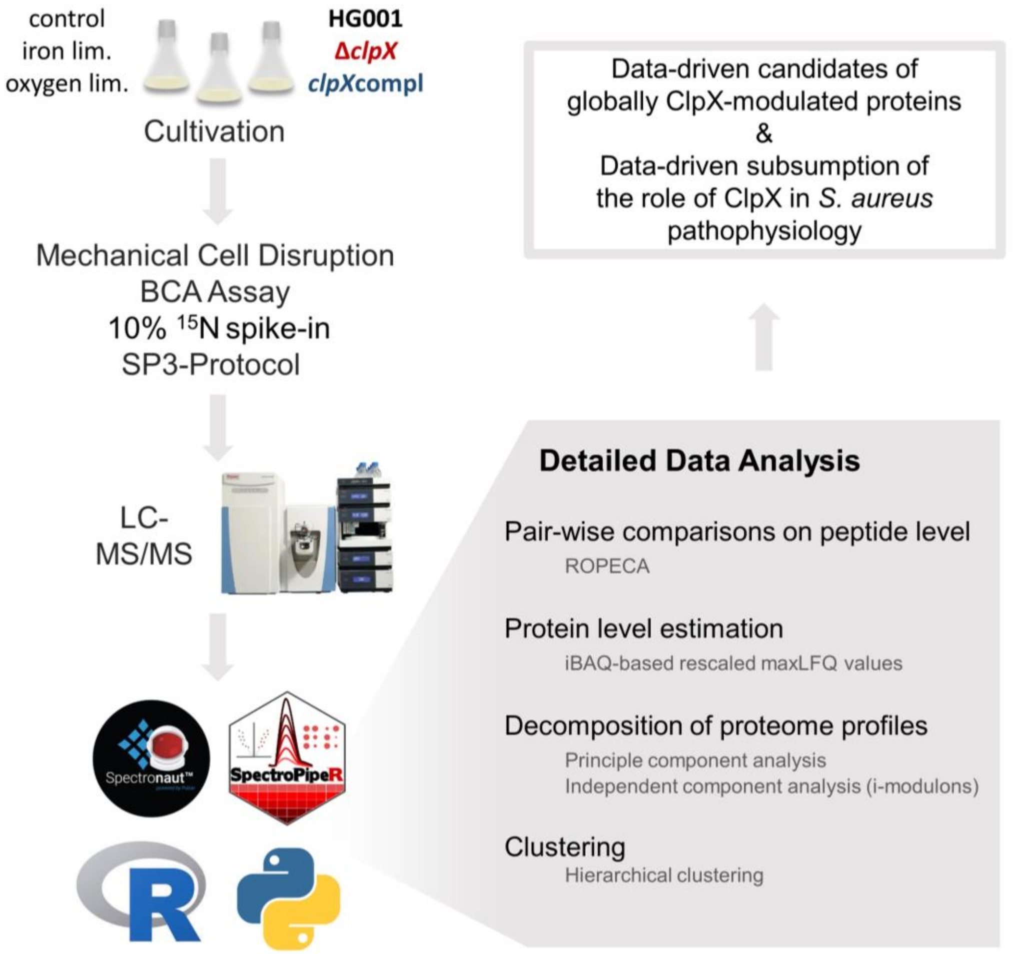

**Figure S4. Workflow of the conducted global label-free proteome analysis.** ClpX-proficient and -deficient *S. aureus* strains HG001, Δ*clpX* and *clpX*compl were cultivated under control, iron limitation and oxygen limitation conditions. Samples were harvested during the exponential and stationary growth phase and the cellular proteome was recorded *via* LC-MS/MS in DIA mode. Proteome data was normalized using an external standard ($^{15}N$ spike-in) for reliable quantification of the in protein composition-varying samples. Detailed data analysis was conducted to enable the identification of new candidates of ClpX-modulated proteins and to provide a resource to gain insights into the role of ClpX in *S. aureus* pathophysiology.

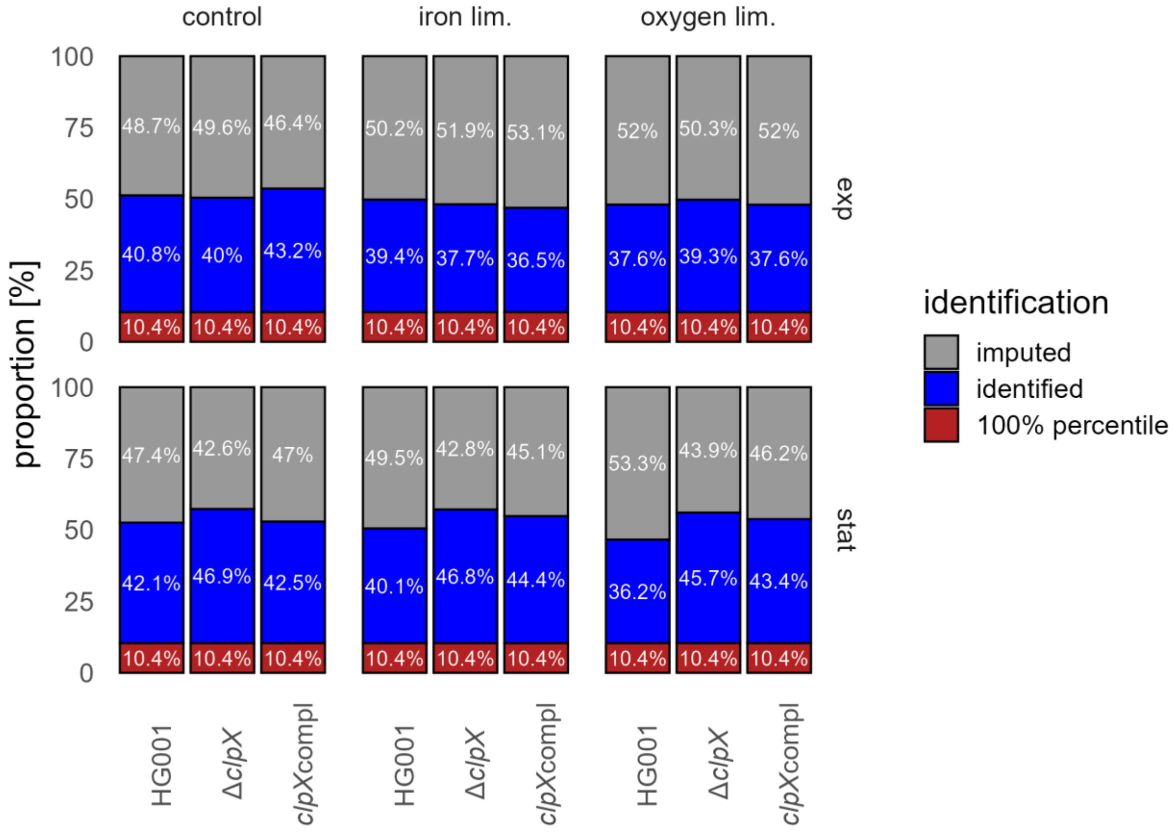

**Figure S5. Peptide ion composition across the experimental conditions of the proteome data set.** Identification numbers of ions were calculated for each sample. Mean proportion of each experimental condition is depicted. 100% percentile: Ions identified in all samples. Identified: Ions identified with q-value ≤ 0.001. Imputed: Profiled ions and ions with q-value > 0.001.

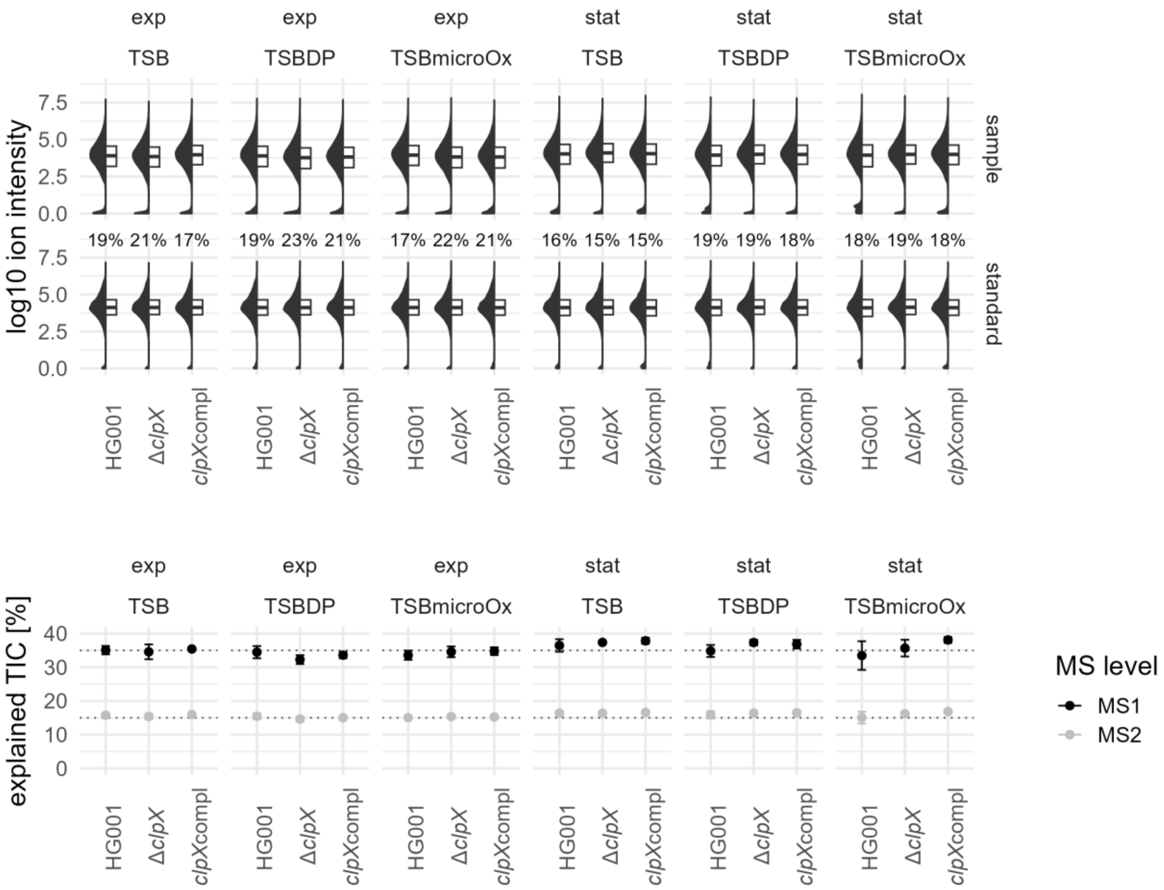

**Figure S6. Overview of peptide ion intensity post normalization and explained total ion currents.** Upper track: $log_{10}$ ion intensity of the sample ions and the external standard ions summarized for the four biological replicates per experimental condition. Percentage numbers indicate the proportion of standard intensity of the explained ions per experimental condition. Lower track: Explained total ion current at MS1 and MS2 level according to the Spectronaut software per experimental condition. TSB: control condition; TSBDP: iron limitation; TSBmicroOx: oxygen limitation.

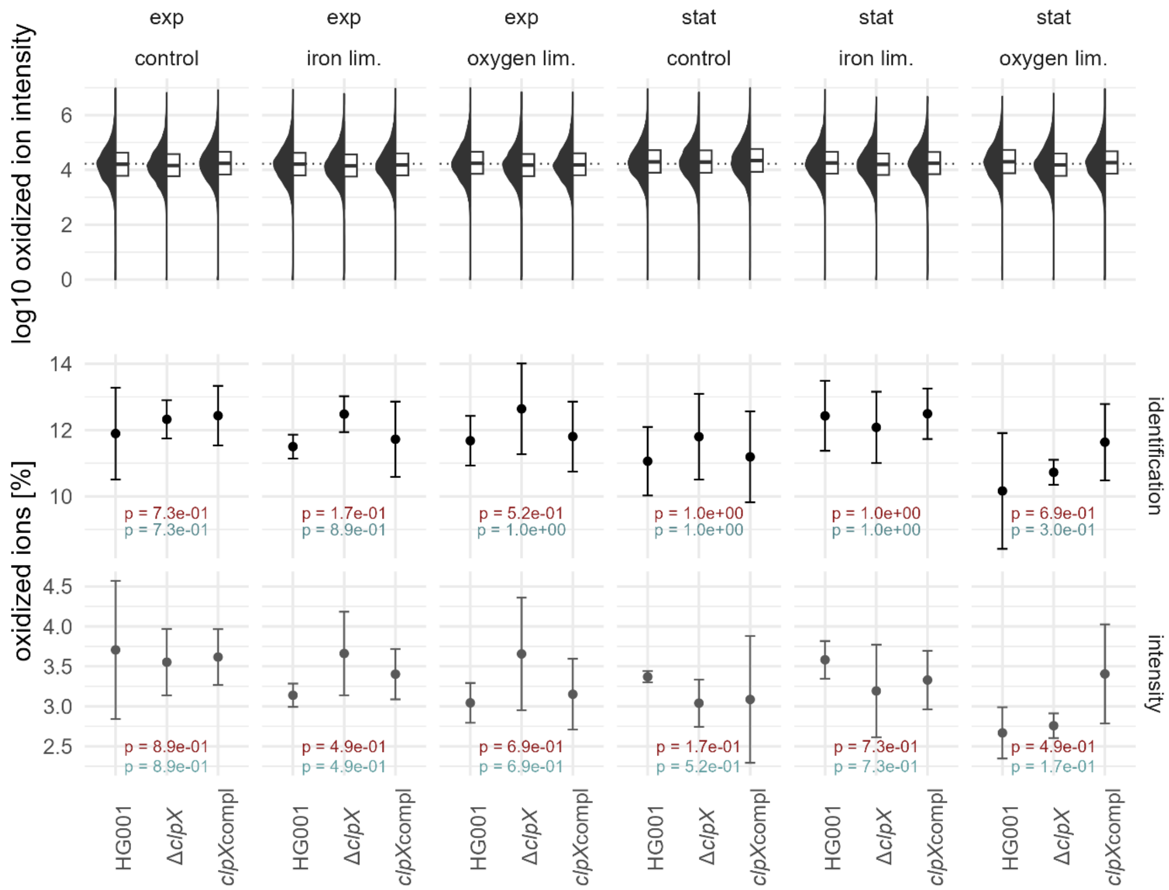

**Figure S7. Quantification of methionine oxidized peptides.** Upper track: log10 ion intensities of methionine oxidized peptide ions across the experimental conditions. Dotted line indicate the global median. Lower track: Proportion of identified (peptide ion q-value ≤ 0.001) oxidized peptides at identification and intensity level. P-values: FDR-adjusted p-values of Wilcoxon test comparing the proportions of Δ*clpX* mutant to the HG001 wild type (red) and the *clpX* complementant to the HG001 wild type (blue).

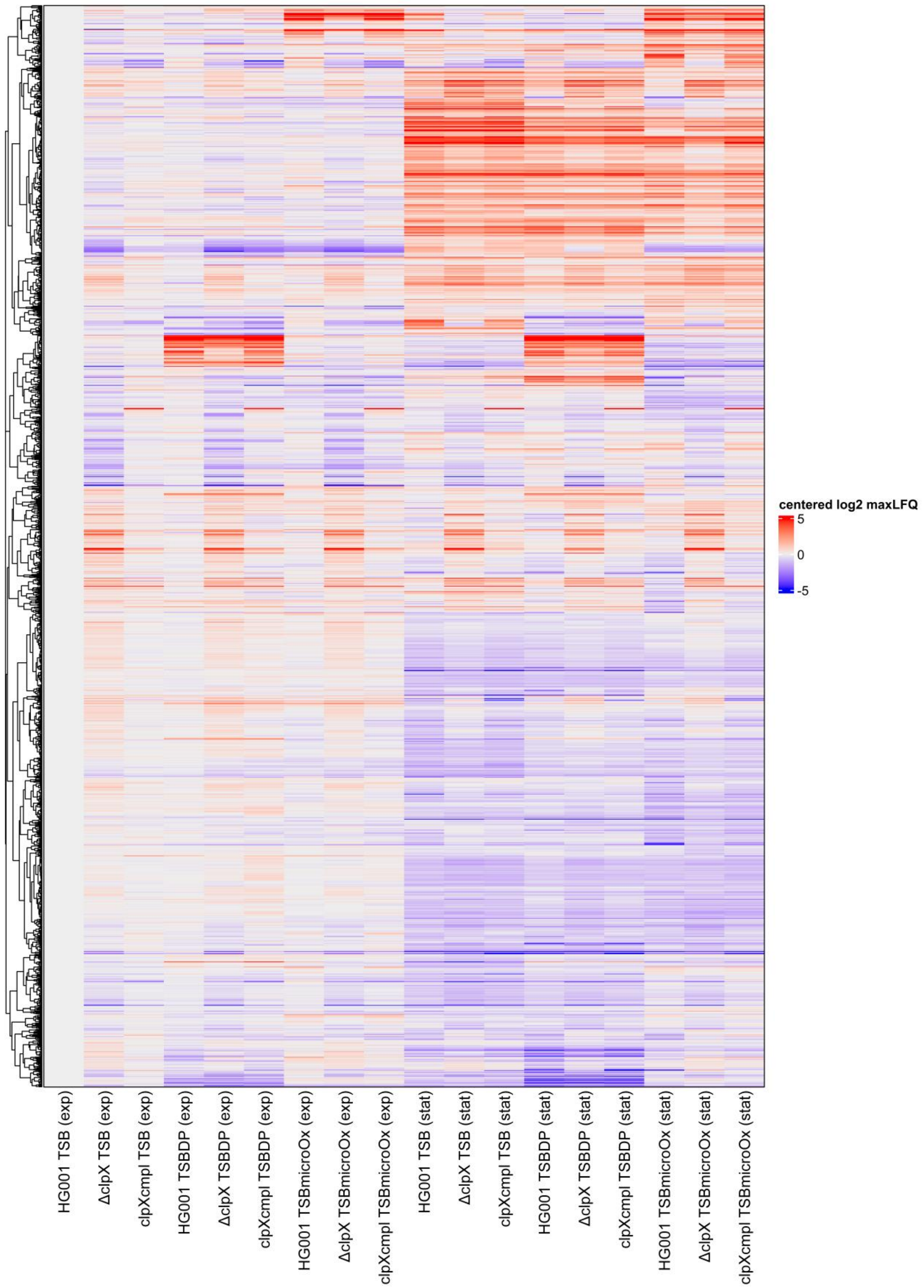

**Figure S8. Global overview of centered log$_2$-transformed protein levels.** The maxLFQ protein level estimations scaled to the relative iBAQ values as used for the PCA and ICA were visualized. For each experimental condition, the mean protein level is depicted. Clustering was performed based on Pearson correlation of the protein levels.

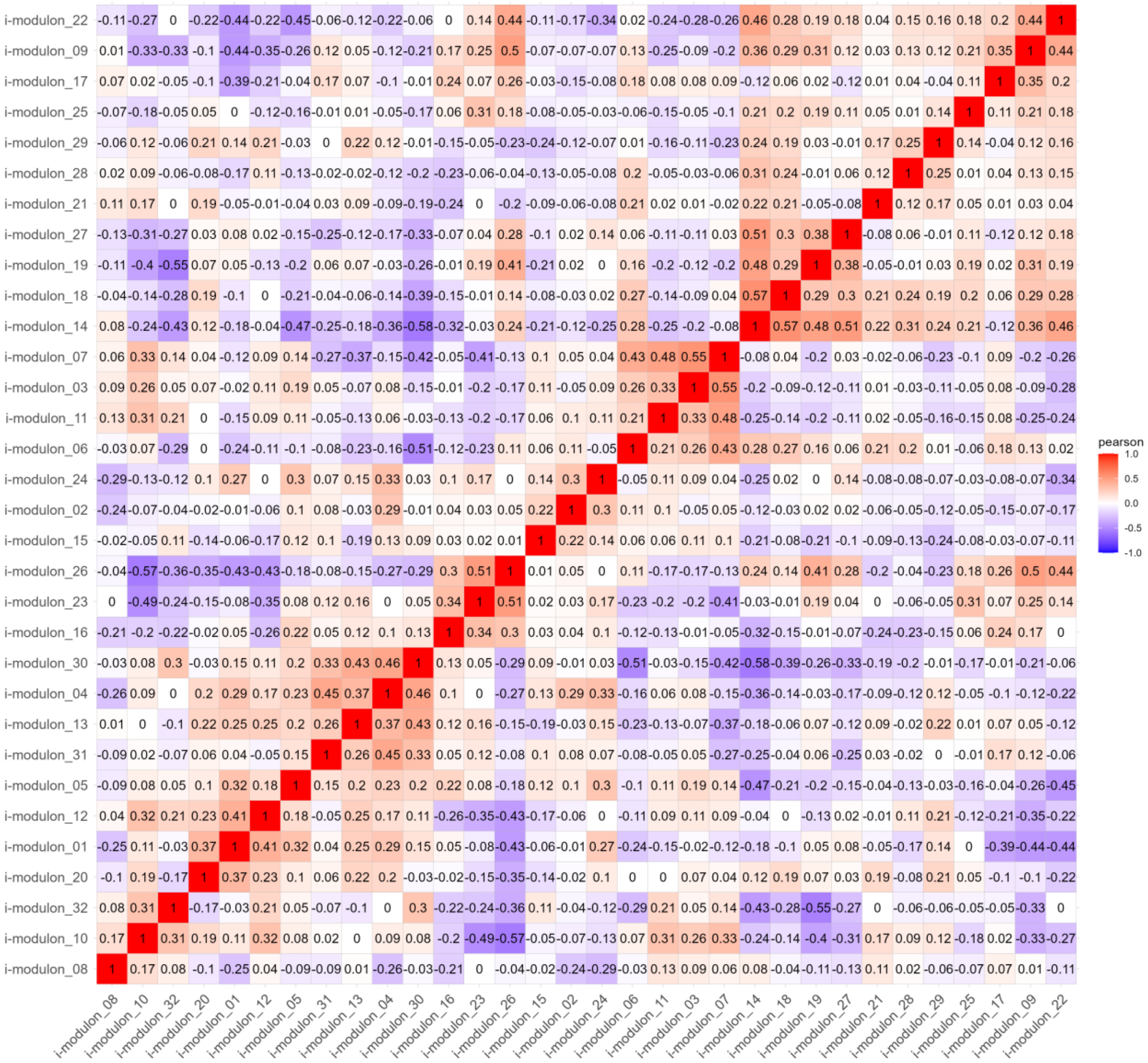

**Figure S9.** **Correlation of i-modulon activities.** The calculated i-modulon activities were correlated using the Pearson method to identify similar i-modulons.

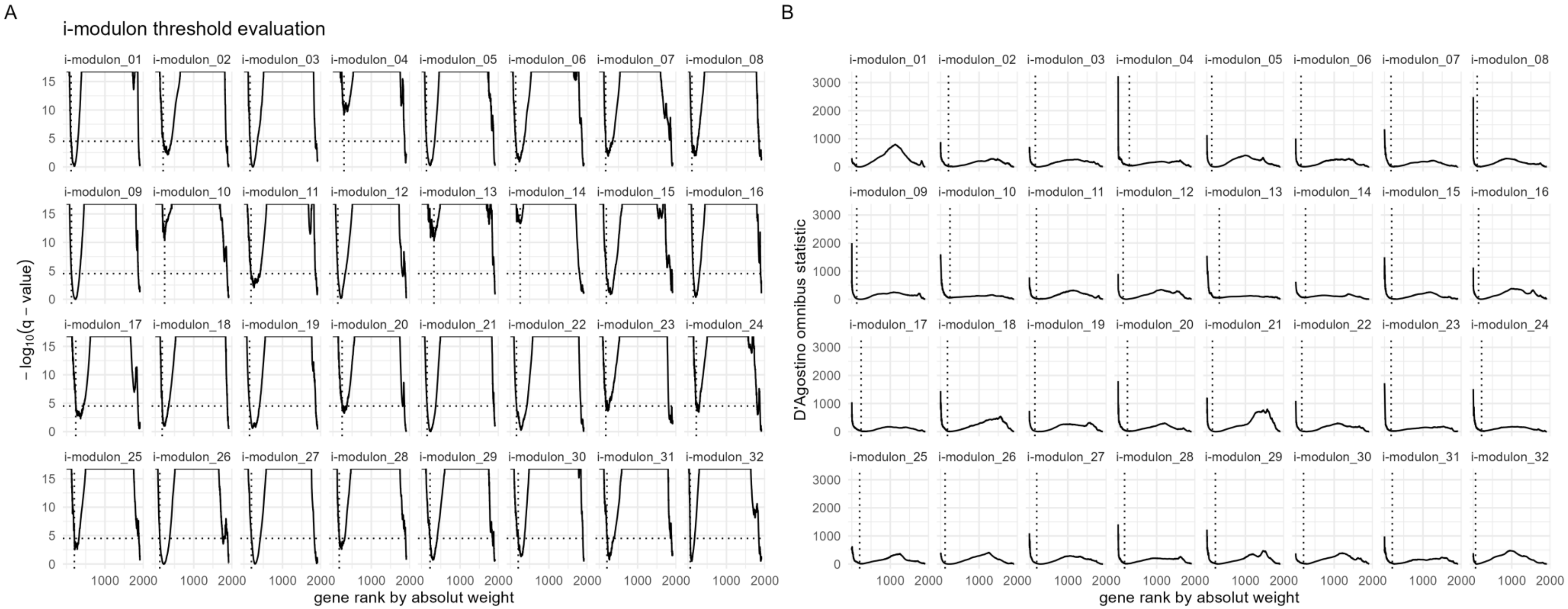

**Figure S10. Evaluation of threshold to define i-modulon member groups.** (A) Overview of the D'Agostino omnibus p-value for investigation of Gaussian-distribution of remaining proteins per i-modulon. Proteins were ranked according to their absolute weight/loading and excluded one by one from the i-modulon. P-values were Bonferroni-adjusted for the 32 i-modulons. Q-value cut-off was set to 0.01 or if not reached in the first half of the protein list to the maximum of the first half of the protein list. Dotted horizontal line indicates the 0.01 threshold. Dotted vertical line depicts the selected cut-off rank. (B) Overview of the D'Agostino omnibus statistic of remaining proteins per i-modulon. Dotted horizontal line indicates the 0.01 threshold. Dotted vertical line depicts the selected cut-off rank.

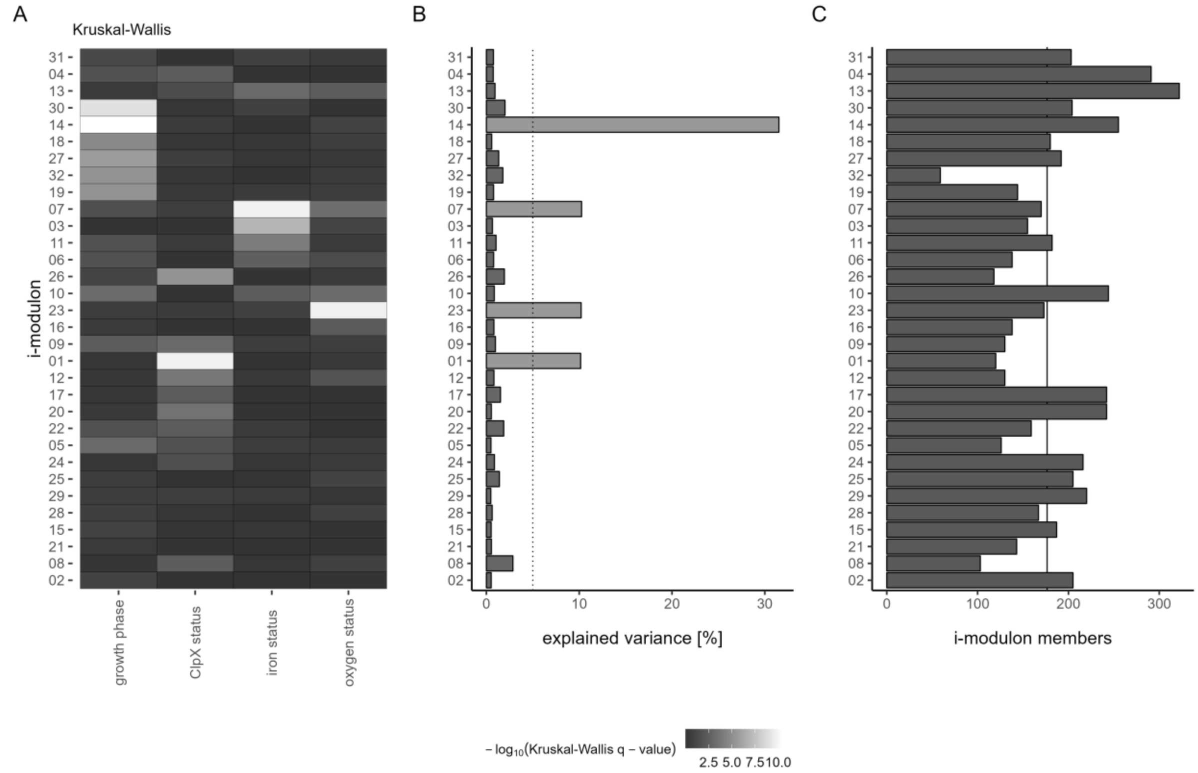

**Figure S11. Additional general aspects of the i-modulons.** (A) Association of i-modulons with experimental conditions. The i-modulon activities were tested for differences according to the experimental conditions growth phase, ClpX status, iron status and oxygen status using the Kruskal-Wallis test. P-values were adjusted using the Benjamini-Hochberg method and association with q ≤ 0.05 were considered significant. The heat map represents the $-\log_{10}$-transformed q-values. (B) Explained variance for each single i-modulon. The dotted line indicates the 5% threshold. I-modulons explaining more than 5% are colored in light grey. (C) Numbers of i-modulon members. For each i-modulon, member proteins were determined based on their weights and the overall non-Gaussian distribution of protein loadings per i-modulon (Figure S10). The median number of i-modulon members is indicated as line.

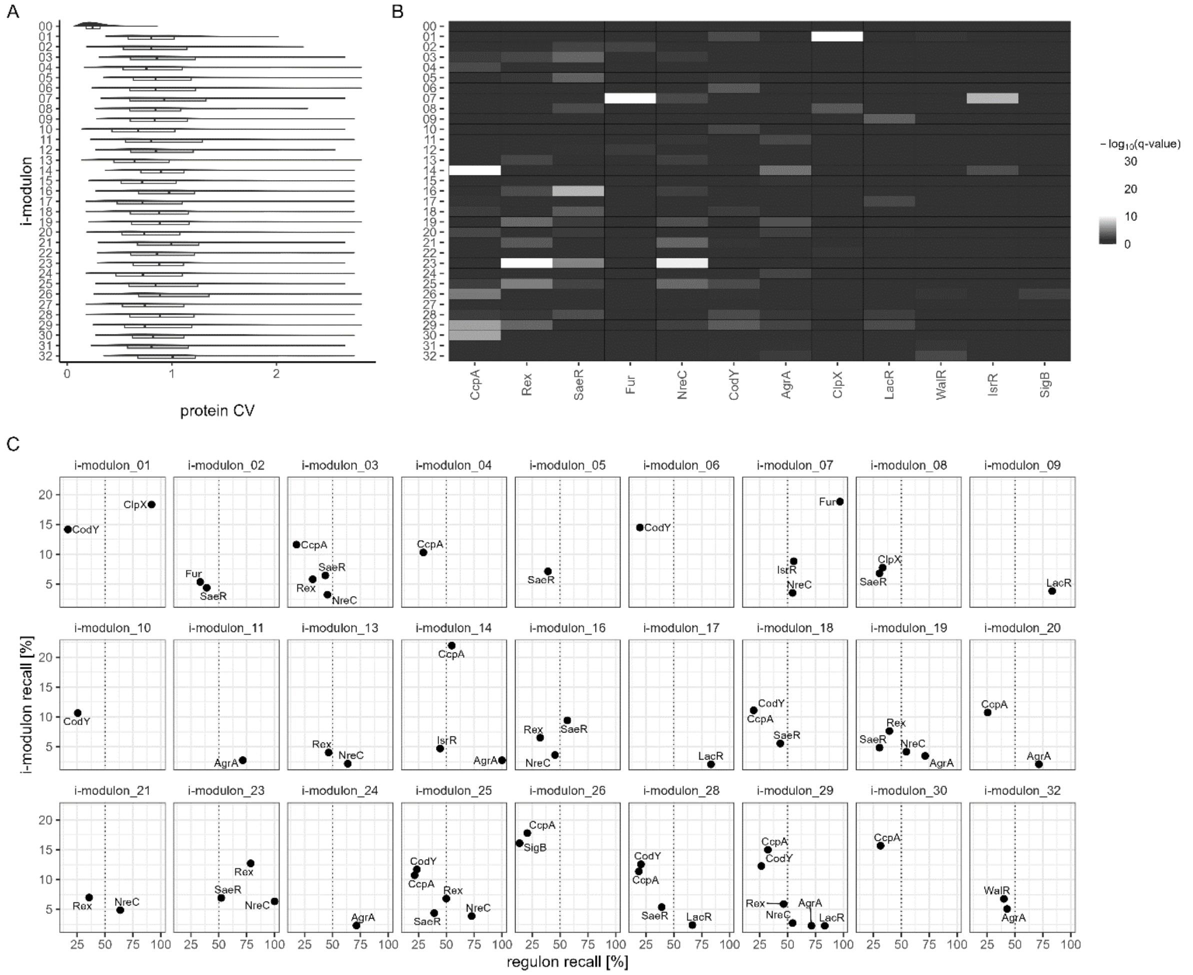

**Figure S12. Investigation of the selected i-modulon member groups.** (A) Comparison of the coefficient of variance of the proteins assigned to the 33 i-modulons. I-modulon_00 represents the group of stable proteins. (B) Overrepresentation analysis of the i-modulon member groups. Overrepresentation was tested using the Fisher's Exact test with Benjamini-Hochberg adjustment of the p-values. Known regulatory entities were depicted with q-value ≤ 0.01 in at least one i-modulon group. Known regulatory entities with at least five detected members were considered. (C) Evaluation of representation regulatory entity groups of i-modulons. Recall of the regulatory group is depicted on the x-axis and recall of the i-modulon is depicted on the y-axis. Dotted line represents 50% recall. Significantly (q-value ≤ 0.01) overrepresented regulatory groups are depicted.

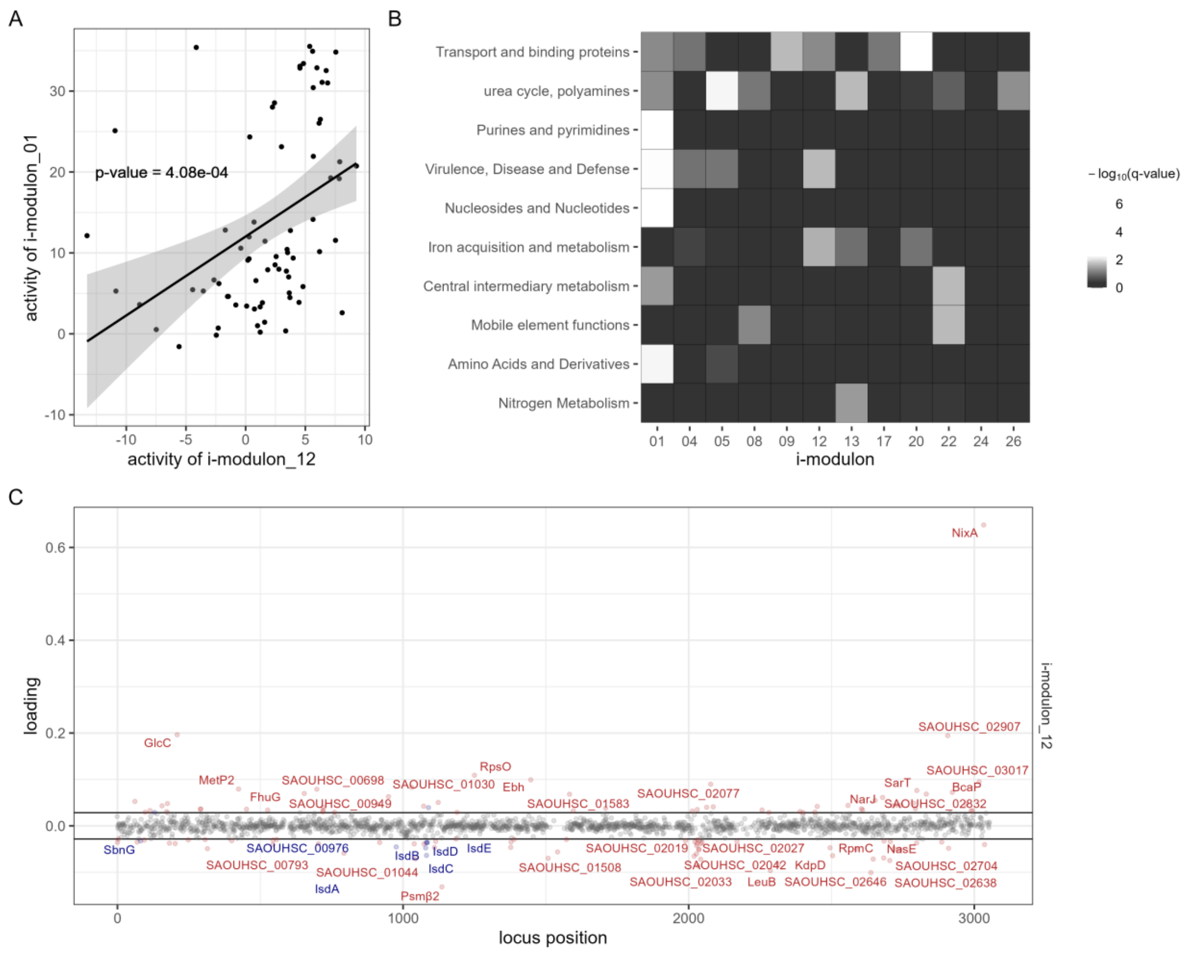

**Figure S13. Additional aspects of ClpX-associated i-modulons.** (A) Correlation of i-modulon activities of the main ClpX-associated i-modulon and i-modulon_12. P-value of Pearson correlation between the activity profiles is indicated. (B) Heat map of overrepresentation analysis of functional annotations for the sub-group of ClpX status associated i-modulons (Figure S11A). TIGRFAM (main role) and TheSEED categories according to *Aureo*Wiki were used as functional categories. Overrepresentation was tested using the Fisher's Exact test with Benjamini-Hochberg adjustment of the p-values. **(C)** Representation of loadings of proteins for the i-modulon_12. Protein loadings were plotted to the genome position. Dotted lines indicate the loading threshold for i-modulon members (Figure S10). I-modulon members are indicated in red and labelled if abs(loading) ≥ 2*threshold. Members categorized as "iron acquisition" function are labelled in blue.

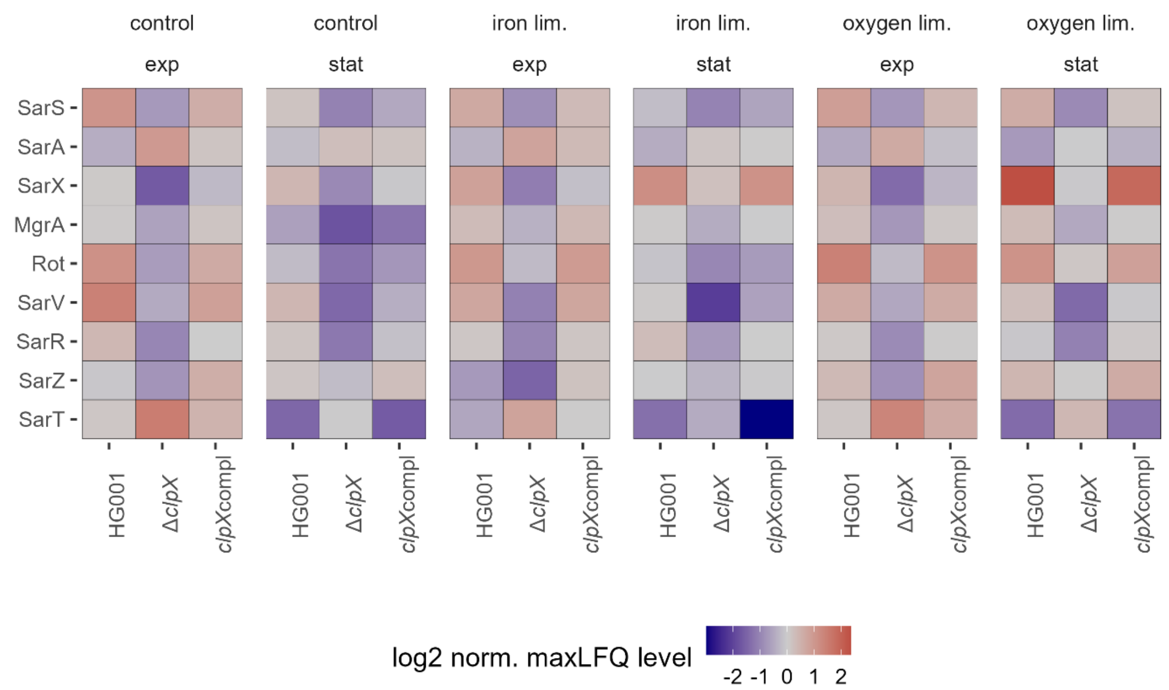

**Figure S14. Heat map of Sar family proteins.** Mean maxLFQ values were centered to the median and log2 transformed.

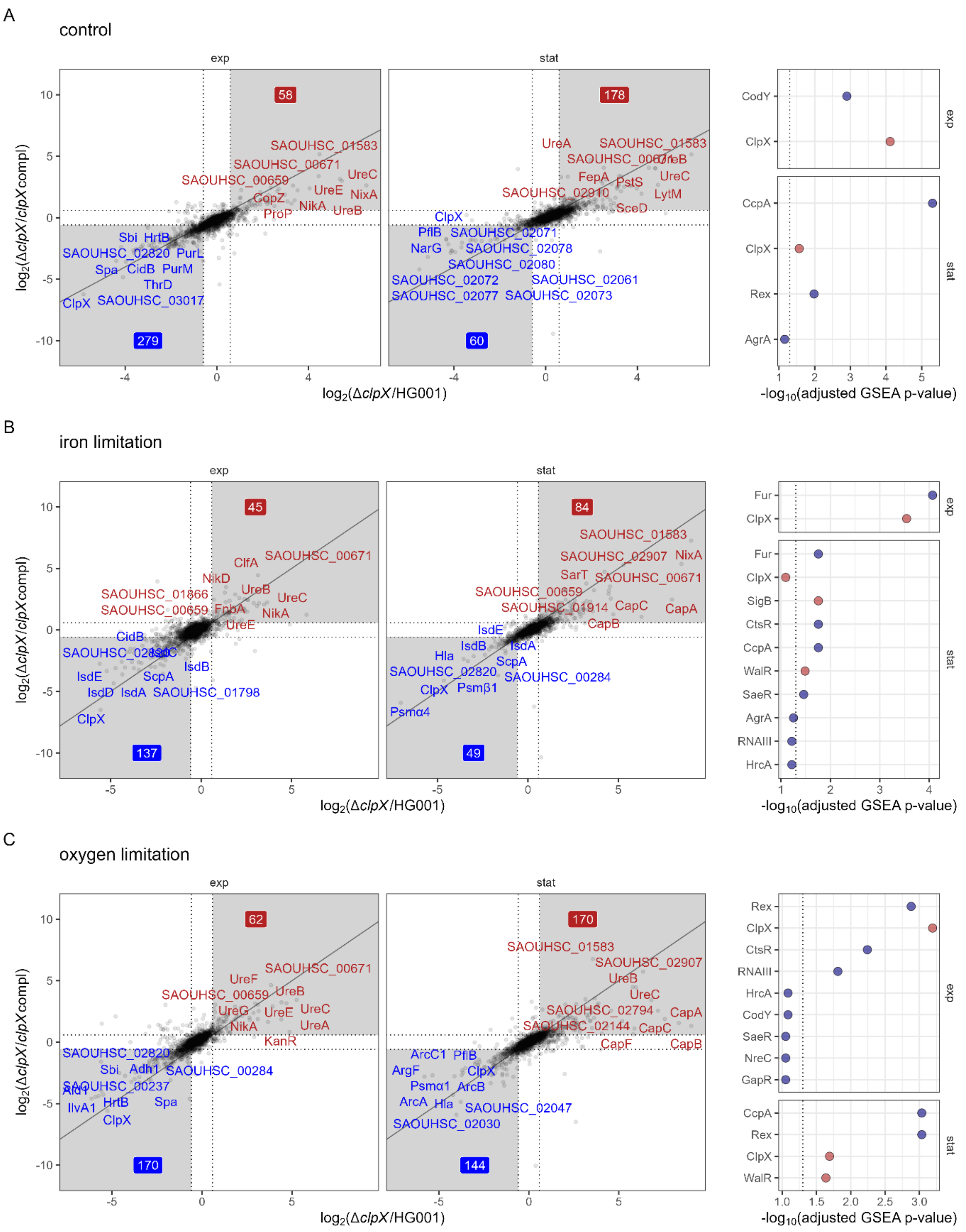

**Figure S15. Investigation of the selected i-modulon member groups.** ClpX modulated proteins under specific stress conditions. ROPECA pairwise statistics were used. Proteins altered in the Δ*clpX* mutant compared to the HG001 wild type and the complemented strain were visualized. Top ten proteins, which were significantly altered (|fold change| ≥ 1.5 & q-value ≤ 0.05) in both comparisons were labelled. The fold change thresholds are indicated as dotted lines. Significantly reduced protein levels are indicated in blue and significantly elevated protein levels are indicated in red. Numbers indicate the total number of significantly reduced and elevated proteins. Comparisons for the exponential and stationary growth phase are separated. GSEA statistics for the analysis are visualized in the right panel. Positive enrichment scores are indicated in red and negative scores are indicated in blue. The dotted line indicates the 0.05 threshold. Proteins were ranked according to the signed Euclidean distance to the zero point considering the $log_2$ ratios and q-values. (A) Control condition. (B) Iron limitation. (C) Oxygen limitation.

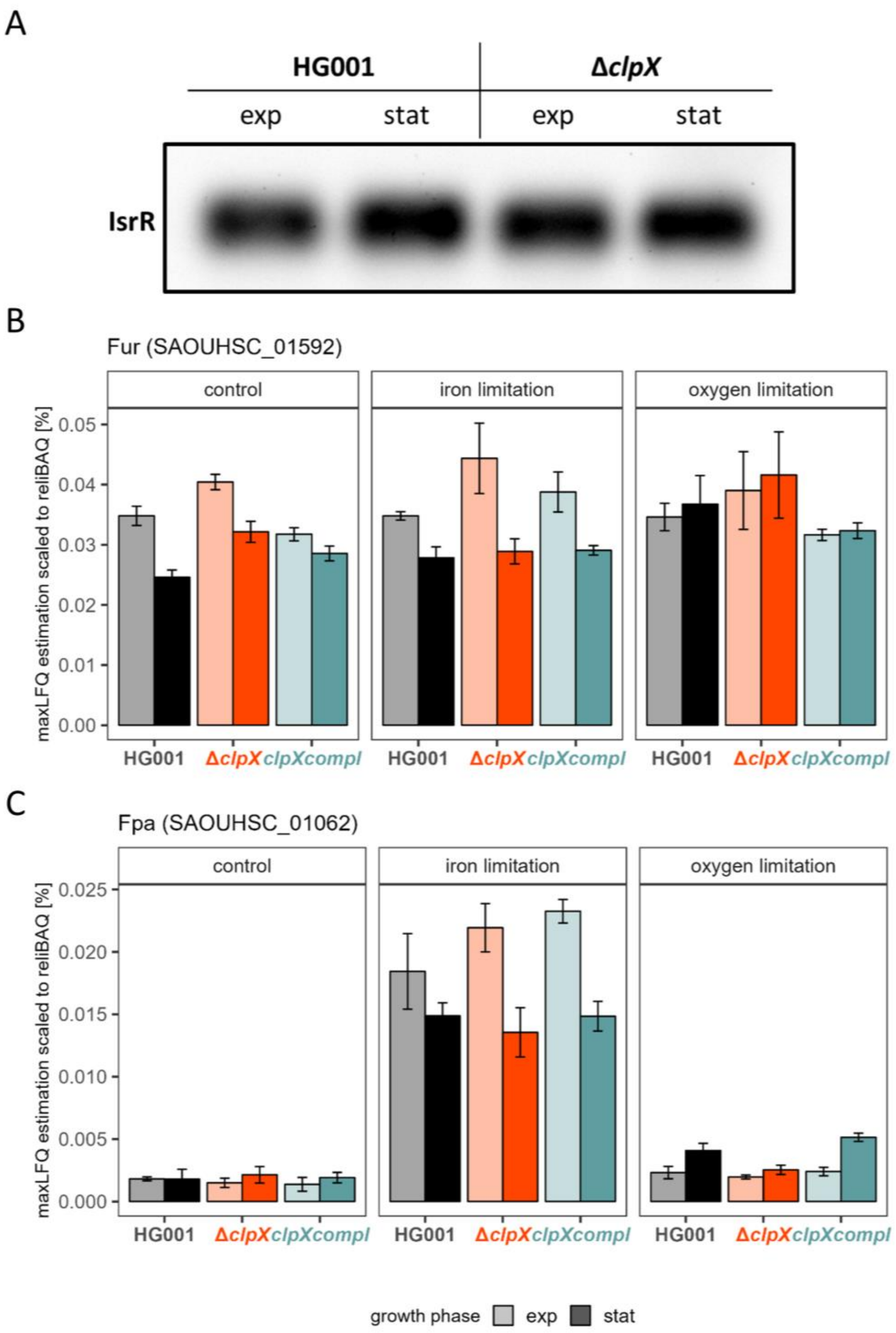

**Figure S16. Overview of known regulators of the iron limitation response.** (A) Northern Blot analysis of IsrR levels in TSBDP during the exponential (2.5h) and stationary (8h) growth phase in the HG001 wild type and the Δ*clpX* mutant. 4 µg of the total RNA was used for each sample. (B) Protein levels of the master regulator Fur. The relative iBAQ-rescaled maxLFQ values are shown. (C) Protein levels of the Fur antagonist Fpa. The relative iBAQ-rescaled maxLFQ values are shown.

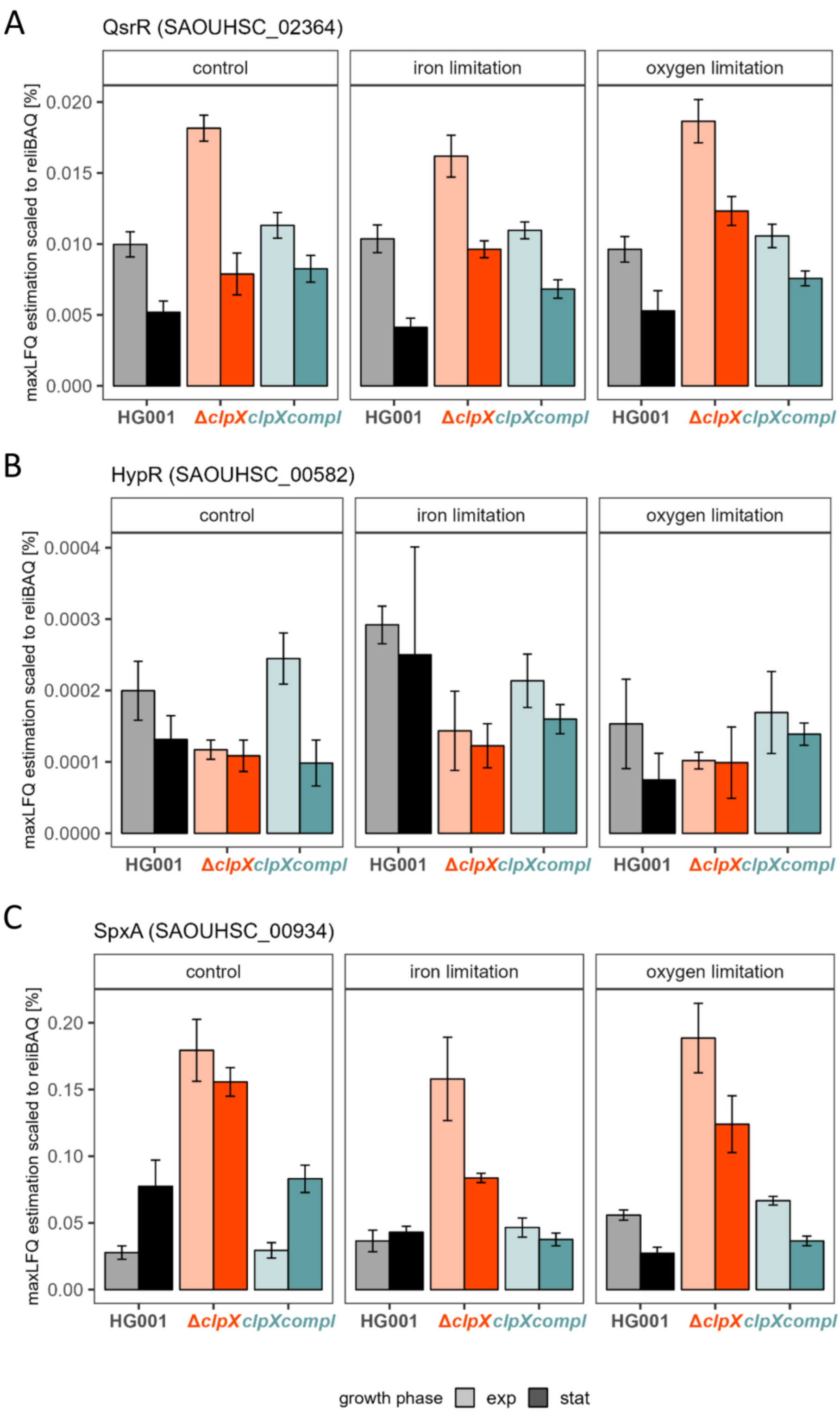

**Figure S17. Overview of regulators of the thiol stress response.** (A) Protein levels of QsrR. (B) Protein levels of HypR. (C) Protein levels of SpxA. The relative iBAQ-rescaled maxLFQ values are shown.

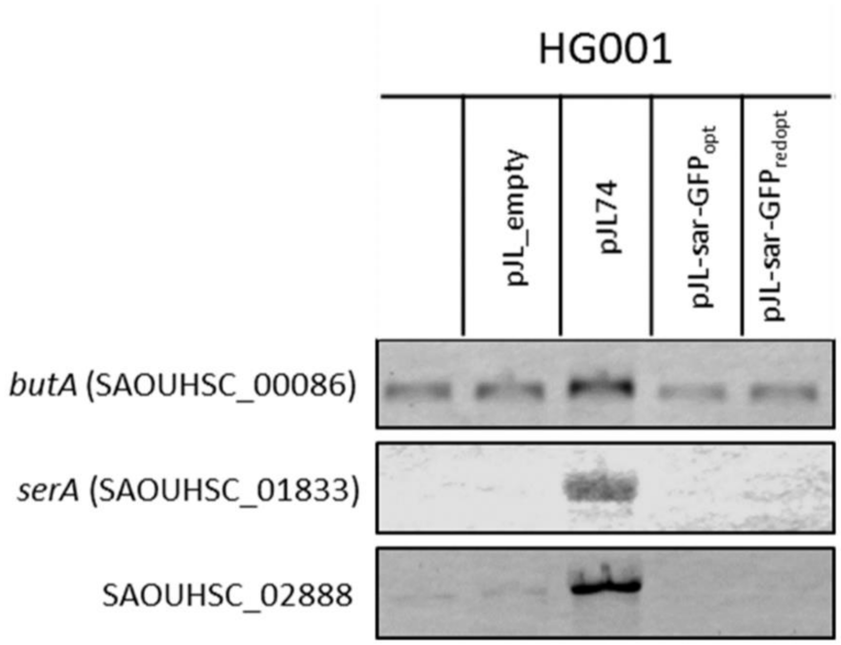

**Figure S18. Induction of the CodY regulon by heterologous expression of non-codon optimized *gfp*.** HG001 carrying an empty pJL control plasmid, the pJL74 plasmid, a pJL74 derivate with codon optimization based on the *S. aureus* codon usage frequency of ribosomal proteins (Michalik *et al.*, 2017) and the pJL-sar-$GFP_{redopt}$ plasmid. Strains were grown in pMEM as indicated in the main manuscript and bacterial cells were harvested at exponential growth phase ($OD_{600nm}$ 0.4). RNA preparation and Northern blotting were performed as described in the main manuscript but using the LI-COR system (LI-COR Biosciences, USA) as described in (Harms *et al.*, 2024). 3 µg of total RNA were used each.

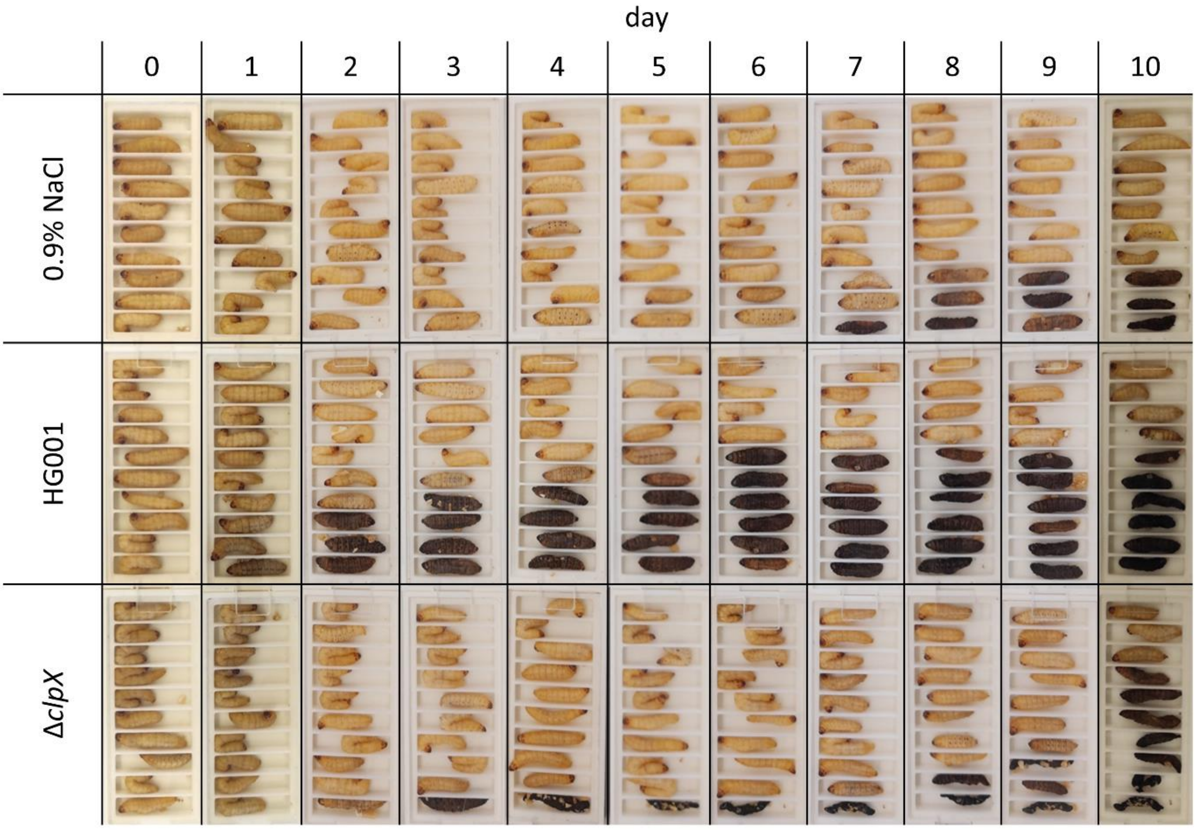

**Figure S19. Representation of the *Galleria mellonella* infection experiment.** *G. mellonella* larvae were infected with 1×$10^5$ *S. aureus* cells in 10 µl 0.9% NaCl solution or as treatment control 10 µl sterile 0.9% NaCl solution was injected. Per bacterial strain (HG001 wild type and Δ*clpX* mutant) or treatment control and biological independent experiment, ten larvae were infected. In total three biological replicates were performed. One representative biological replicate is shown.

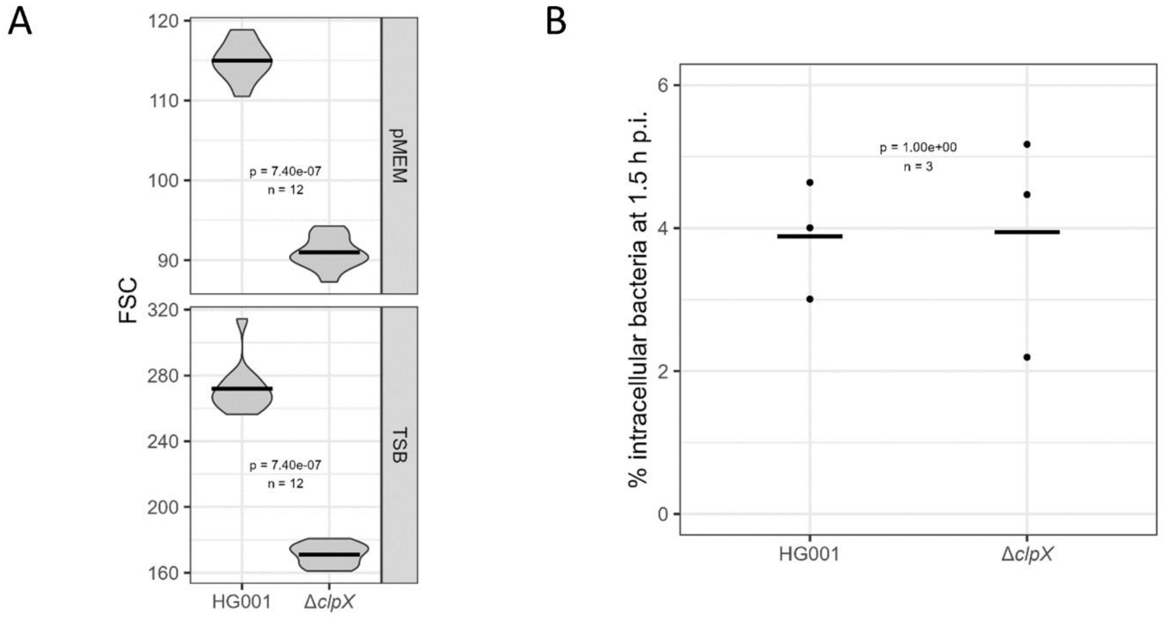

**Figure S20. Comparison of bacterial cell sizes and internalization efficacy of HG001 and Δ*clpX*.** (A) Differences in bacterial cell sizes were estimated *via* measurement of the forward scatter (FSC). HG001 and Δ*clpX* carrying the pJL-sar-$GFP_{redopt}$ plasmid were cultivated in TSB and pMEM to mid-exponential growth phase (TSB: $OD_{600}$ 1; pMEM: $OD_{600}$ 0.4). Two biological replicates with each two technical replicates and each three measurements were obtained. Significance of the FSC difference were tested using the Wilcoxon test. (B) Efficacy of internalization of HG001 and Δ*clpX* by 16HBE14o- cells. Efficacy was estimated by calculation of the proportion of intracellular *S. aureus* cells compared to the whole population at 1.5 h p.i. Three biological replicates were generated. Significance of the difference was evaluated using the Wilcoxon test.

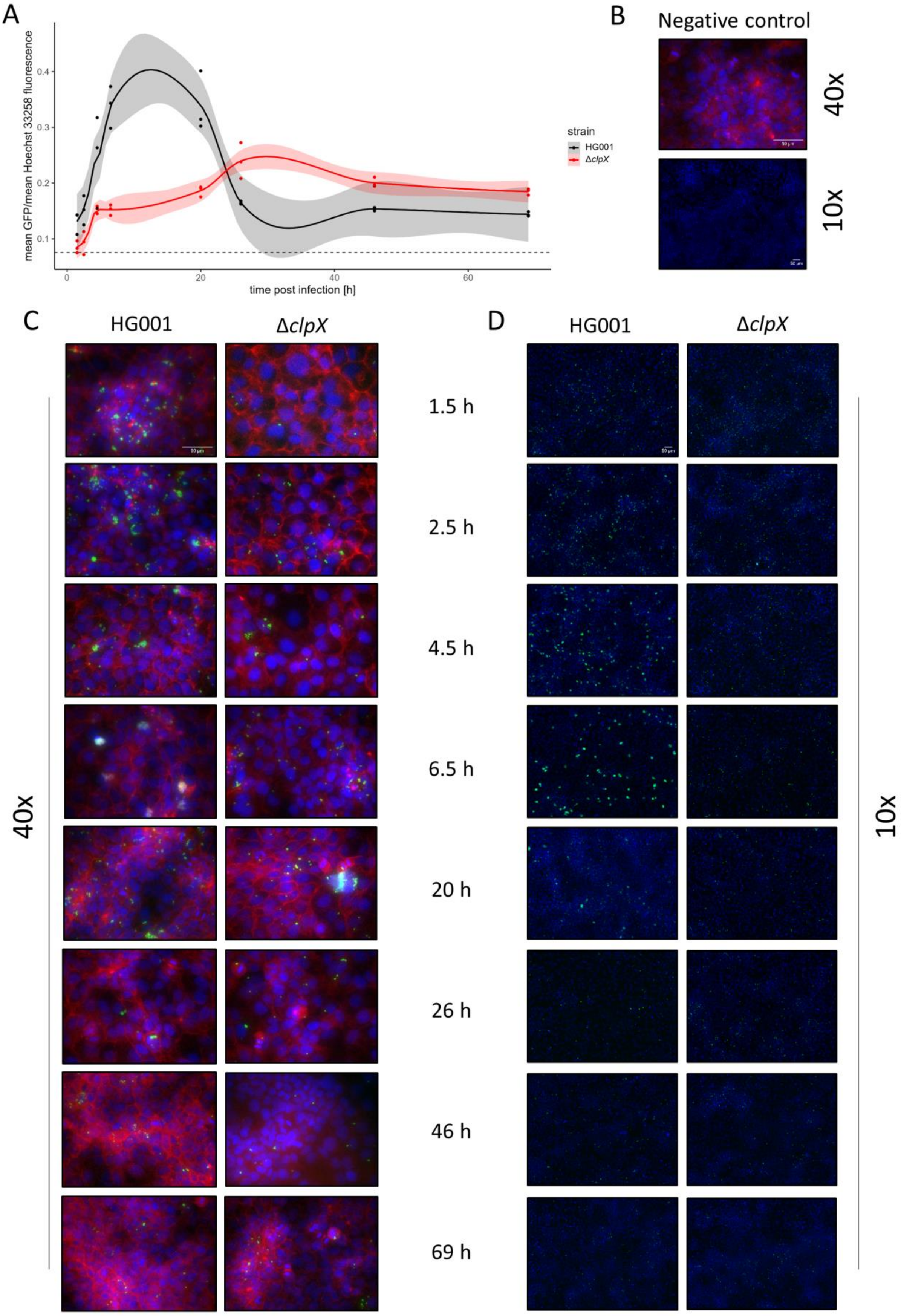
A
mean GFP/mean Hoechst 33258 fluorescence
time post infection [h]
strain
HG001
ΔclpX
B
Negative control
40x
10x
C
HG001
ΔclpX
1.5 h
2.5 h
4.5 h
6.5 h
20 h
26 h
46 h
69 h
40x
D
HG001
ΔclpX
10x

**Figure S21. Microscopic evaluation of the long-term intracellular infection of the 16HBE14o- cells.** Cells were infected with HG001 and Δ*clpX* carrying the pJL-sar-GFP$_{redopt}$ plasmid at a MOI of 60. (A) Quantification of the mean GFP intensity across the images obtained with the 10x objective normalized to the mean Hoechst 33258 fluorescence. The GFP signal represents intracellular *S. aureus* cells and the Hoechst 33258 represent the nuclei of 16HBE14o- host cells. For each time point p.i. three separate regions of interest were randomly chosen. The loess fit of all replicates across the time series per bacterial strain was depicted in bold with the corresponding 95% CI. Mean ratio of the uninfected negative control is depicted as dotted line. (B-D) Representative overlay microscopy pictures using the 10x and 40x objective. GFP signals are displayed in green and Hoechst 33258 signals in blue. At 40x, Phalloidin conjugated Alexafluor 568 was additionally visualized in red. Each channel was separately auto-scaled for visualization purposes. (B) Negative controls. (C) Time series at 40x. (D) Time series at 10x as used for quantification.
